# Long-read sequencing-based atlas of tissue-specific expression of Drp1 transcript variants

**DOI:** 10.1101/2025.10.06.680828

**Authors:** Feng Yan, Jafar S. Jabbari, Naomi X.Y. Ling, Anne M. Kong, Jia Q Truong, Ayeshah A. Rosdah, Christopher Langendorf, Lifang Zhang, Jarmon G. Lees, Sebastian Bass-Stringer, Danise Ann Onda, Kim Loh, Jessica Holien, Sean Lal, Nadia M. Davidson, Jonathan S. Oakhill, Shiang Y. Lim

## Abstract

Dynamin-related protein 1 (Drp1), encoded by *DNM1L*, is essential for mitochondrial fission, but its functional roles remain unclear due to isoform-specific effects from alternative splicing. Short-read RNA sequencing fails to resolve full-length isoforms involving distant exons, limiting our understanding. Here, we applied targeted long-read sequencing to profile full-length *DNM1L* transcripts in human left ventricle and iPSC-derived cardiomyocytes, recovering all annotated isoforms with conserved expression patterns and isoforms 1-4 being most abundant. Functional assays revealed that isoform abundance does not predict enzymatic activity. Extending this to six mouse tissues, we identified distinct, tissue-enriched expression profiles. Functional rescue in Drp1-knockout mouse embryonic fibroblasts showed isoform-dependent differences in mitochondrial fission. Isoforms lacking the A-insert (e.g., b and d) robustly rescued fission, while isoforms enriched in brain or muscle showed only partial rescue, suggesting exons 2 and 3 negatively regulate Drp1 activity. Our cross-species atlas integrates long-read transcriptomics with functional validation, revealing how isoform diversity underpins tissue-specific mitochondrial dynamics and physiological roles of Drp1.

**Summary:** Using long-read sequencing, we mapped full-length *DNM1L/Dnm1l* isoforms in human and mouse tissues, uncovering tissue-specific expression and isoform-dependent mitochondrial fission activity. This reveals how alternative splicing shapes Drp1 function, with implications for understanding its role in health and disease.

## Introduction

Mitochondria are essential organelles within eukaryotic cells, functioning as the primary source of energy production through oxidative phosphorylation. Their morphology is highly dynamic, constantly undergoing processes of fusion and fission [1, 2]. The balance between mitochondrial fusion and fission is crucial for maintaining cellular homeostasis, adapting to various physiological conditions, and responding to cellular stress [3]. Disruption of this balance, particularly through mutations or dysregulation of Dynamin-related protein 1 (Drp1), a key driver of mitochondrial fission, results in fragmented, dysfunctional mitochondria that can lead to cellular damage or death. Drp1 also plays a pivotal role in essential cellular processes such as proliferation, survival, autophagy, and pluripotency, with its absence being embryonically lethal [1, 2, 3, 4]. Despite its importance, the exact mechanisms through which Drp1 influences mitochondrial health remain poorly understood, constituting a significant knowledge gap in mitochondrial biology.

An emerging area of research focuses on the isoforms of Drp1, with eight distinct isoforms identified in humans. These isoforms primarily arise from alternative splicing of exon 3 (encoding the GTPase domain), and exons 16 and 17 (encoding the variable domain) of the *DNM1L* gene [1] (**Figure 1**, **Table 1**). In mice, alternative splicing of exon 1-4 and 16-17 of the mouse *Dnm1l* gene has resulted in the identification of 20 Drp1 isoforms (**Figure 2**, **Table 2**). A study by Strack and colleagues reported tissue-specific expression of certain Drp1 isoforms in mice. For example, isoforms containing exon 3 (equivalent to human isoform 5, 6 and 8) are highly expressed in the brain and skeletal muscle, while those lacking exon 16 but containing exon 17 (equivalent to human isoform 2) are primarily found in the kidney, spleen and heart [5]. Additionally, excessive levels of specific isoforms, such as isoform 5, may be linked to disease pathologies, including brain tumours [6]. Recently, distinct splice variants of Drp1 have been shown to differentially influence cell proliferation, migration, sensitivity to chemotherapeutic agents, and tumour growth *in vivo*. The splice variant containing the full-length variable domain (including exons 16 and 17) plays a more active role in promoting mitochondrial fission. In contrast, the variant lacking exon 16 contributes less significantly to this process [7]. These isoforms display tissue-specific expression profiles, suggesting that they may play distinct roles in cellular functions [1, 6, 8]. However, the incomplete understanding of their expression patterns, with only a few studies reporting differential expression, coupled with the lack of clarity regarding how each isoform contributes to mitochondrial dynamics, complicates our comprehension of the role of Drp1 in health and disease. Moreover, many studies on Drp1 have failed to clearly specify which isoform was investigated or have overly focused on certain isoforms [1, 8, 9, 10], which may not be representative of the full range of Drp1 activities. This lack of specificity in research has contributed to conflicting findings, hindering the development of effective therapeutic strategies targeting Drp1.

**Figure 1.**
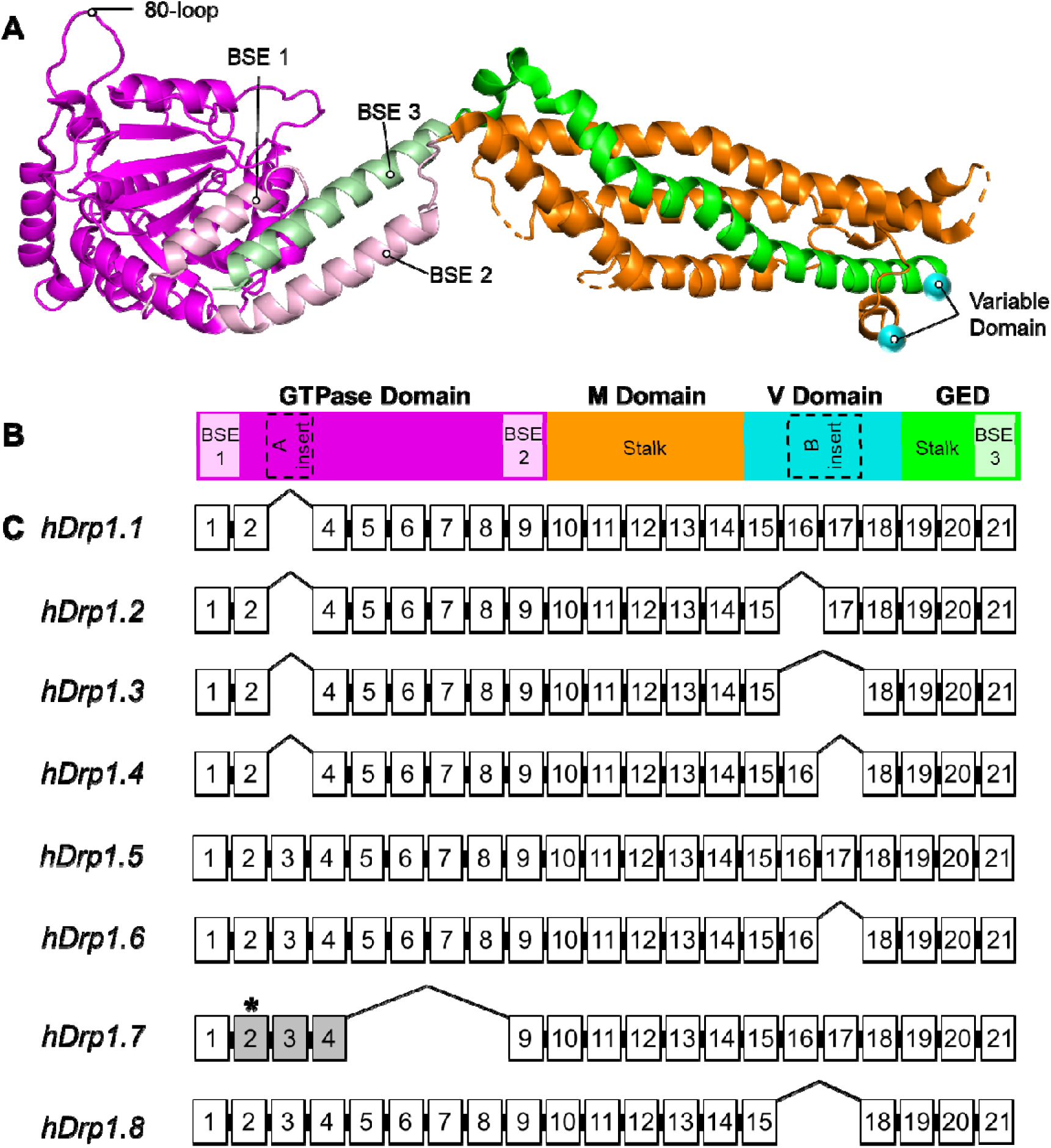
Structure and coding region of human Drp1 isoforms. **(A)** Cartoon representation of the composite Drp1 model assembled from crystal structures (PDB IDs: 4H1V, residues 1–285; and 4BEJ, residues 282–701). The structure is color-coded by domain: the GTPase domain is shown in magenta and includes bundle signalling elements (BSE) 1 and 2 in light pink; the middle (stalk) domain appears in orange; the variable domain is depicted as cyan spheres, representing only the visible N- and C-terminal residues; and the GTPase effector domain (GED) is shown in green, encompassing portions of the stalk and BSE 3 (light green). **(B)** The linear schematic, color-coded to match the structure in panel A, illustrates the domain organisation of Drp1. The A- and B-inserts are depicted as dashed black boxes. **(C)** Coding regions for human Drp1 isoforms 1–8 are depicted with exon numbering based on the coding sequence of isoform 5 (CCDS61095.1). The GTPase domain is encoded by exons 1–9, the middle (M) domain by exons 10–14, the variable (V) domain by exons 15–18, and the GED by exons 19–21. The A-insert, located within exon 3, forms part of the GTPase domain, while the B-insert, spanning exons 16–17, resides within the variable domain. Exons, introns, and domains are not shown to scale. White rectangles represent exons; LJ indicates skipped exons; * marks an alternative start codon; and gray shading highlights regions encoding alternative amino acid sequences.

**Figure 2.**
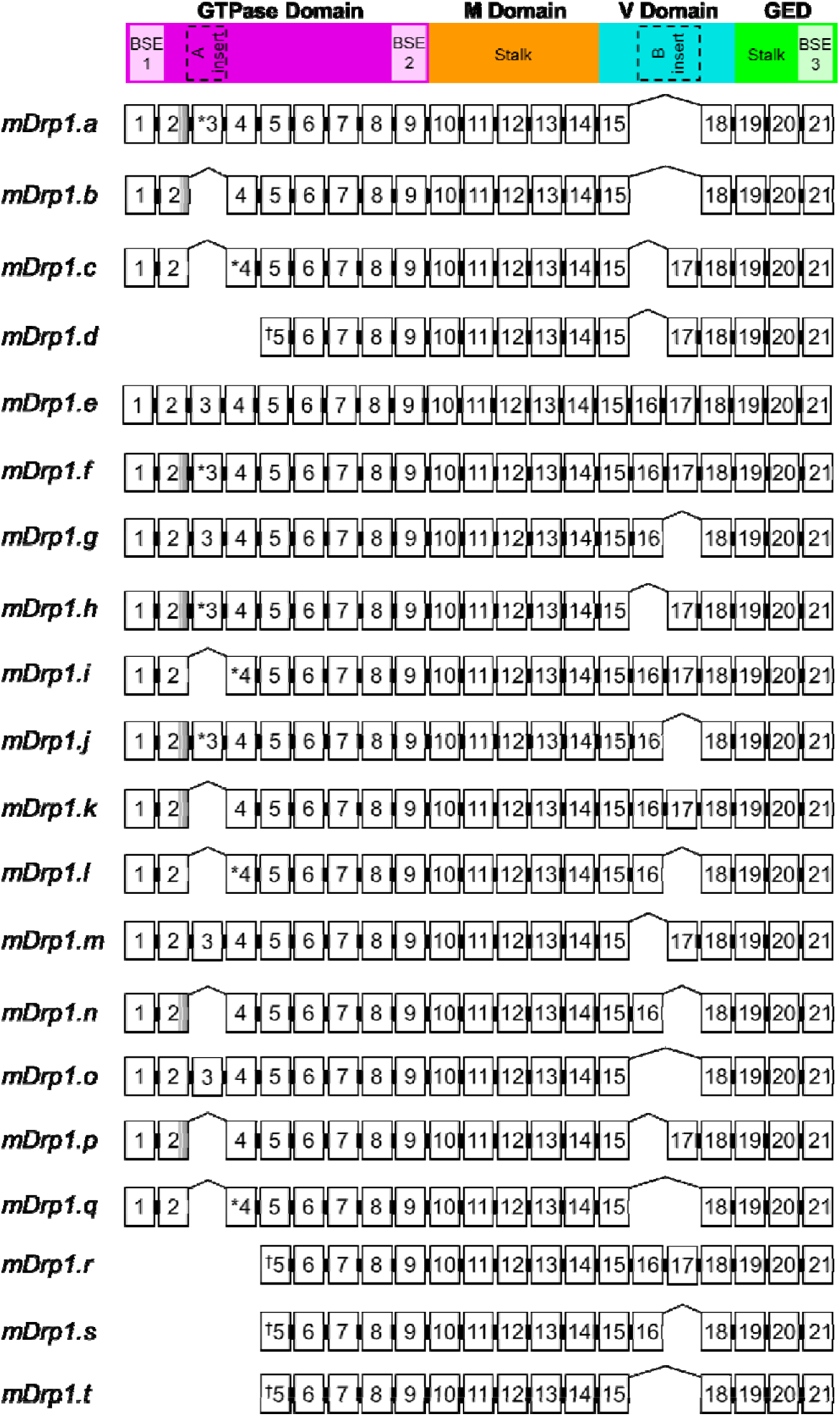
Structure and coding region of mouse Drp1 isoforms. Coding regions of mouse Drp1 isoforms a–t with exon numbering based on the longest isoform, isoform e (mRNA transcript NM_001360007.2, protein NP_001346936.1). The GTPase domain is encoded by exons 1–9, the middle (M) domain by exons 10–14, the variable (V) domain by exons 15–18, and the GTPase effector domain (GED) by exons 19–21. The A-insert, located within exon 3, forms part of the GTPase domain, while the B-insert, spanning exons 16–17, resides within the variable domain. Exons, introns, and domains are not shown to scale. White rectangles represent exons. Squares with shaded grey rectangles on exon 2 denote missing amino acid compared to mDrp1.e (GKFQSW) at C-terminus. * on exon 3 and 4 denote first amino acid is different from isoform e (N>D for exon 3) and (G>R for exon 4). ^†^ on exon 5 denote addition of amino acid (M) at N-terminus.

**Table 1:** Human DRP1 isoforms and *DNM1L* splice variants.

| **Isoform ID** | **Transcript Accession** | **Transcript Length (nucleotides)** | **Transcript Protein Accession** | **Transcript Protein Length (amino acids)** |
| --- | --- | --- | --- | --- |
| 1 | NM_012062.5 | 4514 | NP_036192.2 | 736 |
| 2 | NM_012063.4 | 4436 | NP_036193.2 | 710 |
| 3 | NM_005690.5 | 4403 | NP_005681.2 | 699 |
| 4 | NM_001278463.2 | 4481 | NP_001265392.1 | 725 |
| 5 | NM_001278464.2 | 4553 | NP_001265393.1 | 749 |
| 6 | NM_001278465.2 | 4520 | NP_001265394.1 | 738 |
| 7 | NM_001278466.2 | 4110 | NP_001265395.1 | 533 |
| 8 | NM_001330380.2 | 4442 | NP_001317309.1 | 712 |

**Table 2:** Mouse Drp1 isoforms and *Dnm1l* splice variants.

| **Isoform ID** | **Transcript Accession** | **Transcript Length (nucleotides)** | **Transcript Protein Accession** | **Transcript Protein Length (amino acids)** |
| --- | --- | --- | --- | --- |
| a | NM_152816.4 | 4023 | NP_690029.2 | 712 |
| b | NM_001025947.3 | 3984 | NP_001021118.1 | 699 |
| c | NM_001276340.2 | 4035 | NP_001263269.1 | 716 |
| d* | NM_001405252.1 | 3970 | NP_001392181.1 | 612 |
| d* | NM_001276341.2 | 3790 | NP_001263270.1 | 612 |
| e | NM_001360007.2 | 4152 | NP_001346936.1 | 755 |
| f | NM_001360008.2 | 4134 | NP_001346937.1 | 749 |
| g | NM_001360009.2 | 4119 | NP_001346938.1 | 744 |
| h | NM_001360010.2 | 4056 | NP_001346939.1 | 723 |
| i | NM_001405253.1 | 4113 | NP_001392182.1 | 742 |
| j | NM_001405254.1 | 4101 | NP_001392183.1 | 738 |
| k | NM_001405255.1 | 4095 | NP_001392184.1 | 736 |
| l | NM_001405256.1 | 4080 | NP_001392185.1 | 731 |
| m | NM_001405257.1 | 4074 | NP_001392186.1 | 729 |
| n | NM_001405258.1 | 4062 | NP_001392187.1 | 725 |
| o | NM_001405259.1 | 4041 | NP_001392188.1 | 718 |
| p | NM_001405260.1 | 4017 | NP_001392189.1 | 710 |
| q | NM_001405261.1 | 4002 | NP_001392190.1 | 705 |
| r | NM_001405262.1 | 4048 | NP_001392191.1 | 638 |
| s | NM_001405263.1 | 4015 | NP_001392192.1 | 627 |
| t<sup>#</sup> | NM_001405264.1 | 3937 | NP_001392193.1 | 601 |
| t<sup>#</sup> | NM_001405265.1 | 3757 | NP_001392194.1 | 601 |

\*NM\_001405252.1 and NM\_001276341.2 encode the same isoform d but differ in their 5' untranslated regions.
<sup>#</sup>NM\_001405264.1 and NM\_001405265.1 encode the same isoform t but differ in their 5' untranslated regions.

The structural differences between Drp1 isoforms, such as variations in protein domains and GTPase activity, may impact their oligomerization and interactions with other cellular proteins [1, 8, 11, 12, 13]. These differences could affect the accessibility of binding sites for small molecule inhibitors or peptides, influencing their potential as therapeutic agents. Some studies indicate that certain isoforms may be more involved in regulating mitochondrial fission, apoptosis, or other cell functions in specific tissues. For example, isoform 2 has been shown to confer resistance to apoptosis in some models, whereas isoforms containing specific exons, such as isoforms 1 and 6, may interact with anti-apoptotic proteins like Bcl-xL [5, 14]. These findings suggest that selectively targeting Drp1 isoforms could open new avenues for disease-specific treatments, particularly in conditions like neurodegenerative diseases, cardiovascular disorders, and cancer.

Given the complexity of Drp1 isoforms generated through alternative splicing and their potential contribution to cellular dysfunction, it is critical to investigate their tissue-specific expression, structural diversity, and functional consequences. Short-read sequencing has limited full-length isoform characterization, obscuring their biological significance. In this study, we applied long-read transcriptomics together with functional assays to comprehensively define *DNM1L/Dnm1l* isoforms, revealing distinct tissue-enriched expression patterns and isoform-dependent differences in mitochondrial fission activity. By establishing a cross-tissue atlas of *DNM1L/Dnm1l* expression and function in mouse, we provide new insights into how Drp1 isoform diversity shapes mitochondrial dynamics, with implications for developing more precise therapeutic strategies for mitochondrial-related diseases.

## Methods

### Animal tissues

Animal tissue sample collection and experimental procedures were approved by the Animal Ethics Committee of St. Vincent’s Hospital (AEC 027/22; Victoria, Australia) and were conducted in accordance with the Australian National Health and Medical Research Council guidelines for the care and maintenance of animals. Adult male C57BL/6 mice (8 weeks old) were euthanised via cervical dislocation. Tissues and organs were promptly harvested and snap-frozen in liquid nitrogen. Total RNA was extracted using the Aurum™ Total RNA Fatty and Fibrous Tissue Kit (BioRad). Briefly, tissues were immersed in 1 mL of RNAlater™ Stabilization Solution (ThermoFisher) and placed on ice. They were then cut into 3 mm^3^ pieces and mechanically homogenized using a TissueRuptor (Qiagen) for 30-60 seconds in PureZOL lysis reagent. The homogenate was incubated at room temperature for 5 minutes and subsequently centrifuged at 12,000 g for 10 minutes at 4°C to pellet insoluble debris. After phase separation, the aqueous phase was processed through a silica spin column. On-column DNase I digestion was performed to remove genomic DNA. Total RNA was eluted in 40□µL of RNase-free water and stored at -80°C.

### Human heart tissues

Donor hearts were obtained from the Sydney Heart Bank, which is approved by the Ethics Committee of The University of Sydney (USYD #2021/122). The hearts were deemed suitable for heart transplantation but unable to be transplanted for reasons including transportation logistics, immune incompatibility, and donor-recipient mismatch in size. They were procured as previously described [15, 16, 17]. The hearts underwent formal pathological examination and were deemed structurally normal. Left ventricle (LV) samples were snap frozen in liquid nitrogen (-196 °C) within 15 mins of harvest. Approximately 40 mg of frozen heart tissues (LV samples from male donors aged 21–56-year-old) were transferred to a 2 mL tube containing one 5 mm stainless steel bead (Qiagen) and was disrupted and homogenised in TRIzol reagent (Invitrogen) using the TissueLyser LT (Qiagen). RNA extraction was then performed according to TRizol manufacturer’s instructions. The extracted RNA was then DNase treated using the Qiagen RNAse-Free DNase Set and subsequently purified using the RNase Mini kit (Qiagen) according to manufacturer’s instructions. The concentration and quality of the RNA were assessed using a Nanodrop and the integrity of the RNA was assessed using an RNA Nano Chip on an Agilent Bioanalyzer.

### Cell culture

The human iPS-Foreskin-2 (CL2) [18] and CERA007c6 (CERA) [19] induced pluripotent stem cells (iPSCs) were maintained on vitronectin-coated plates in TeSR-E8 medium (STEMCELL Technologies, Vancouver, Canada) according to the manufacturer’s protocol. Human cardiomyocytes were derived from iPSCs using a previously described method [20, 21]. Briefly, iPSCs were seeded onto hESC-qualified Matrigel (Corning) coated plates at a density of 1.25×10^5^cells/cm^2^ in TeSR-E8 medium supplemented with 10 µM Y-27632 (Abcam). After 48 hours, medium was replaced with RPMI 1640 basal medium (Thermo Fisher Scientific) containing B-27 without insulin supplement (Thermo Fisher Scientific), growth factor reduced Matrigel (1:60 dilution; Corning), and 10 µM CHIR99021 (STEMCELL Technologies) (designated as day 0). After 24 hours, medium was replaced with RPMI 1640 basal medium containing B-27 without insulin supplement for 24 hours. On day 2, medium was replaced with RPMI 1640 basal medium containing B-27 without insulin supplement and 5 µM IWP2 (Tocris Bioscience) for 72 hours. From day 5 onwards, cells were cultured in RPMI 1640 basal medium containing B-27 supplement (Thermo Fisher Scientific) and 20 ng/mL L-ascorbic acid 2-phosphate sesquimagnesium salt hydrate (Sigma-Aldrich) (referred to as CM medium), and the medium was refreshed every 2-3 days. On day 12, cardiomyocytes were dissociated into single cells and split at a 1:4 ratio onto hESC-qualified Matrigel coated plates in DMEM/F-12 GlutaMAX media (Thermo Fisher Scientific) supplemented with 20% foetal bovine serum (Sigma-Aldrich), 100 µM 2-mercaptoethanol (Thermo Fisher Scientific), 100 µM non-essential amino acids (Thermo Fisher Scientific), 50 U/mL penicillin/streptomycin (Thermo Fisher Scientific), and 10 µM Y-27632. At day 13, medium was replaced with CM medium. From days 14 to 19, cardiomyocytes were enriched to >90% cardiac troponin T positive cells by culture in glucose-free DMEM medium (Thermo Fisher Scientific) containing 4 mM sodium L-lactate (Sigma-Aldrich). Total RNA was isolated from enriched day 19-23 iPSC-derived cardiomyocytes using the Illustra RNAspin Mini Kit (Cytiva) according to the manufacturer’s instructions. DNase I enzyme was used to remove genomic DNA. RNA was eluted (twice) off the column in 40 µL of water, quantified using a 2000c Thermo Nanodrop (Thermo Scientific) spectrophotometer, and stored at -80°C. RNA quality and integrity were assessed using the Agilent TapeStation system, and only samples with acceptable RNA integrity values were included for downstream analysis.

### cDNA synthesis and Nanopore sequencing

The process of cDNA synthesis and long-read sequencing using Nanopore platform is summarised in Figure 3A. For each sample, 300 ng of high-quality total RNA (RIN>8) were reverse transcribed using Maxima H Minus Reverse Transcriptase (Thermo Fisher Scientific) in a 22.5 μL reaction. The reaction included an oligo(dT) primer NaVNP: 5’- ACTTGCCTGTCGCTCTATCTTCTTTTTTTTTTTTTTTTTTTTVN - 3’, a template-switch oligo with unique molecular identifier 10XTSOUMI: 5’-AAGCAGTGGTATCAACGCAGAGTACANNNYRNNNYRNNNYRNNNVTrGrGrG-3’, and RNaseOUT (ThermoFisher Scientific). Reverse transcription was performed at 42 °C for 90 minutes, followed by enzyme deactivation at 85 °C for 5 minutes. The resulting cDNA was purified using 0.7x SPRIselect beads (Beckman Coulter) and eluted in 16 μL. The cleaned cDNA was amplified using KAPA HiFi HotStart ReadyMix (Roche). Initial denaturation was performed at 95 °C for 3 minutes, followed by 17 cycles of 98 °C for 20 seconds, 62 °C for 15 seconds, 72 °C for 3 minutes and a final extension at 72 °C for 5 minutes. The reaction included biotinylated HEx21-R1: 5’-/5Biosg/AAGATGAGTCTCCCGGATTTCAGC-3’or MEx21-R1: 5’-/5Biosg/AAGATGAGTCTCTCGGATTTCAGCA-3’ primers targeting exon 21 in human and mouse Drp1 transcripts, respectively, and ScDNARP: 5’-AAGCAGTGGTATCAACGCA-3’ for TSO binding. Amplified products were cleaned using 0.45x SPRIselect beads and eluted in 12 μL. Biotinylated amplicons were captured using 5 μL of Dynabeads M270 Streptavidin (ThermoFisher Scientific) according to manufacturer instructions. The beads were resuspended in 12 μL. The captured cDNA was re-amplified with KAPA HiFi HotStart ReadyMix using Ex21-R1 (human): 5’-AAGATGAGTCTCCCGGATTTCAGC-3’ or MEx21-R1-2 (mouse): 5’-AAGATGAGTCTCTCGGATTTCAGCA-3’ and cDNARP: 5’-AAGCAGTGGTATCAACGCAGAG-3’primers.

**Figure 3.**
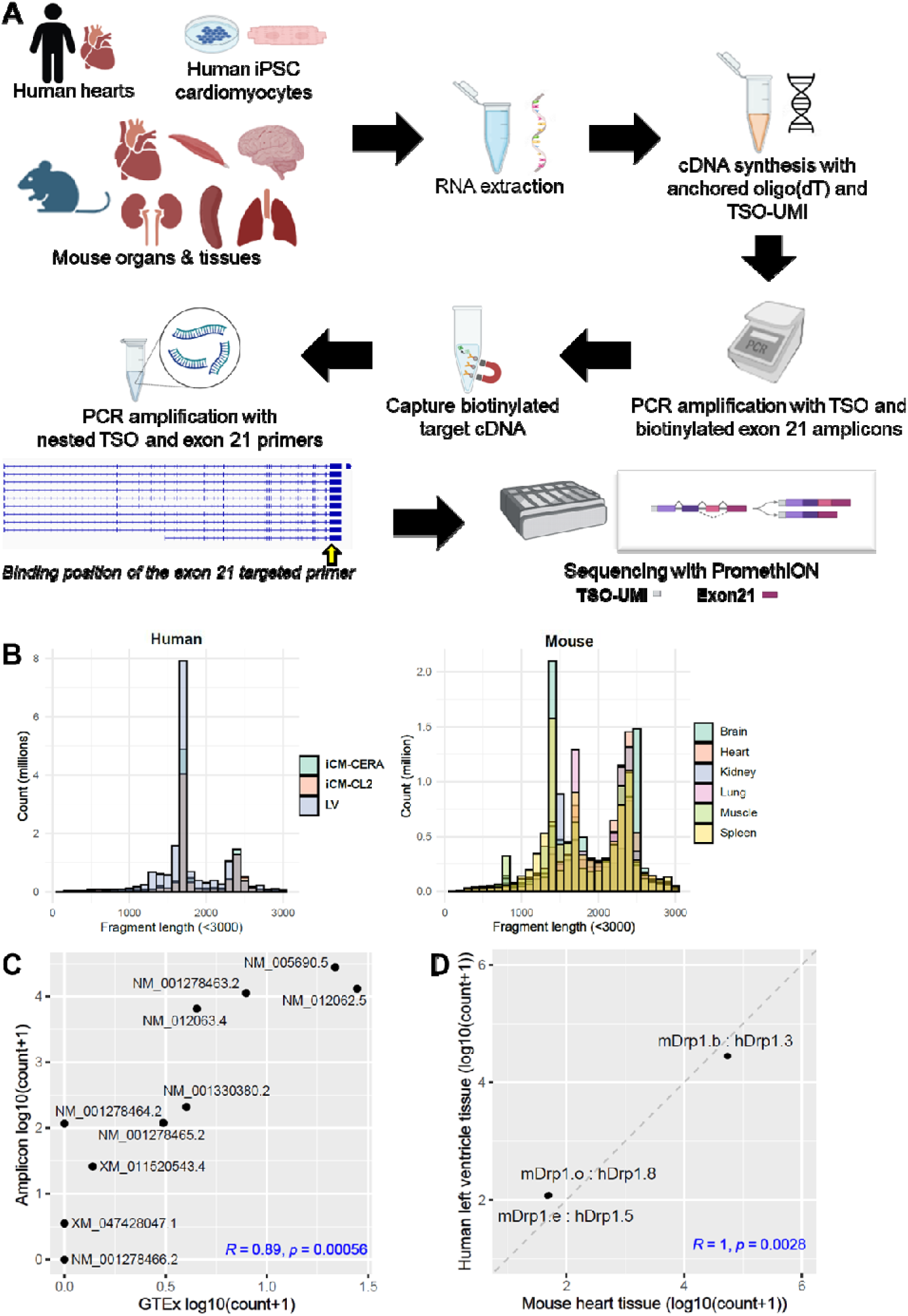
Long-read transcriptomic profiling and cross-species comparison of *DNM1L/Dnm1l* isoforms. **(A)** long-read transcriptomic sequencing workflow. **(B)** Read length distributions across biological human and mouse samples. iCM-CERA (CERA iPSC-derived cardiomyocytes), iCM-CL2 (CL2 iPSC-derived cardiomyocytes), LV (human left ventricle). **(C)** Scatter plot of expression correlation between amplicon and GTEx datasets (*R* = 0.89, *p* = 0.00056). Transcript expression correlation of human *DNM1L* transcripts between amplicon captured in house data and GTEX data. Expression values were averaged from left ventricle samples only in both datasets, and log10 transformed with offset 1. **(D)** Correlation of conserved human and mouse Drp1 isoforms selected for experimental validation. The label is shown as (mouse isoform id : human isoform id). mDrp1.b: hDrp1.3 (NM_001025947.3 : NM_005690.5), mDrp1.e : hDrp1.5 (NM_001360007.2 : NM_001278464.2), mDrp1.o : hDrp1.8 (NM_001405259.1 : NM_001330380.2). Expression was calculated by averaging human left ventricle samples and mouse heart tissues, and log 10 transformed with offset 1.

Thermal cycling conditions were the same as above. Amplified products were cleaned with 0.45x SPRIselect and eluted in 20 μl. Libraries were prepared using the Native Barcoding kit (SQK-NBD114.24, Oxford Nanopore Technologies) and loaded onto a PromethION FLO-PRO114M flow cell for sequencing. Base calling was performed with MinKNOW 23.11.7 super accuracy mode. All oligos were purchased from IDT.

### Bioinformatic analysis of Nanopore targeted sequencing

Basecalled Nanopore FASTQ files were processed with flexiplex (v1.01) [22] to remove barcodes and identify UMI sequences, then mapped to the genome with minimap2 (v2.26-r1175) [23]. For mouse, the GRCm39 genome and NCBI RefSeq annotation (GCF_000001635.27) were used; for human, the GRCh38 genome and NCBI RefSeq annotation (GCA_000001405.15) were used. On target rate was calculated using the reads mapping to Drp1 gene region divided by the total number of mappable reads. UMI-tools (v1.1.6) [24] was used to remove PCR duplicates based on UMIs and genomic coordinates for reads mapped to the Drp1-containing chromosome. Saturation analysis was performed by down sampling reads mapped to the Drp1-containing chromosome and performing UMI-tools deduplication on a subset of the data. Transcript quantification was performed using Bambu (v3.6.0) [25] with UMI-tools deduplicated BAM files and the corresponding annotation. Seven left-ventricle samples were downloaded from GTEx [26], aligned to the human genome, and quantified directly with Bambu.

To generate transcript coverage plots, we extracted reads mapped to the Drp1-containing chromosome and realigned them to the Drp1 cDNA sequences. The FLAMES::get_coverage function (v2.1.9) [27] was run on the realigned BAMs to generate transcript coverage plots. To generate genomic coverage plots, we used deepTools (v3.5.5) [28] on genomic BAM files and summarised the regions using a GTF file containing all exons. BAM files were visualised in IGV to produce representative track views.

The tissue specificity index (TSI) [29] for each transcript was calculated as previously described. Briefly, TSI was calculate using the following formula:

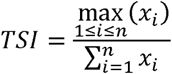

*xi* was the average expression (TPM) in a given tissue, and *n* was the number of tissues. Transcripts were then categorized as tissue-specific (TSI ≥ 0.8), broadly expressed (TSI *<* 0.5), or biased toward a group of tissues (0.5 ≤ TSI *<* 0.8). The scripts for analysis are available on GitHub https://github.com/alexyfyf/Drp1_targeted_ONT.

### Generation of stable Drp1 isoform expression in mouse embryonic fibroblasts (MEFs)

cDNAs encoding for mouse Drp1 isoform b (NM_001025947.3), isoform d (NM_001405252.1), isoform e (NM_001360007.2) and isoform o (NM_001405259.1) were custom synthesised by Gene Universal with C-terminal FLAG-tags and cloned into pLeGO-iG2 using EcoRI/NotI restriction sites. All constructs were sequence-verified. Drp1 wildtype and knockout (KO) MEFs were kindly provided by Professor Hiromi Sesaki (Johns Hopkins University, USA)[30, 31]. Stable expression of Drp1 isoforms, each bearing a COOH-terminal FLAG tag, was achieved in Drp1 KO MEFs via lentiviral transduction using the LeGO-iG2 vector system. Lentiviral particles were produced by co-transfecting HEK293T cells with the envelope plasmid pCMV-VSV-G (Addgene #8454) and the packaging plasmid psPAX2 (Addgene #12260), following a previously described protocol [32]. Drp1 KO MEFs were transduced with 0.45 µm-filtered viral supernatant in the presence of polybrene, and successfully transduced cells were enriched based on EGFP expression using fluorescence-activated cell sorting.

### Western blot analysis

Drp1 expression was analysed in MEFs by Western blotting. Protein was extracted with 1% (v/v) Triton X-100 lysis buffer (50 mM Tris, pH 7.6, 150 mM NaCl, 10% glycerol) supplemented with 1mM EDTA, 1 mM EGTA, 5 mM sodium pyrophosphate, 50 mM sodium fluoride and protease inhibitor cocktail (Roche). Cell lysates were clarified by centrifugation at 14,000 *g* for 15 minutes at 4°C. Supernatant was collected and stored at -80°C. Protein concentration in each sample was quantified using the bicinchoninic acid assay kit (Thermo Fisher Scientific). Protein was reduced and denatured in 5 X sample buffer containing 4mM Tris(2-carboxyethyl)phosphine (TCEP) followed by heating at 90°C for 5 minutes. 40 µg of protein was subjected to 4-20% SDS-PAGE (BioRad Cat#4561094) at 120 V for 60 minutes. Gel was transferred onto a polyvinylidene difluoride membrane (GE Healthcare Life Sciences, Australia) at 70 V for 90 minutes, 4°C, and blocked with Intercept phosphate-buffered saline blocking buffer (LiCOR Biosciences, NE,USA) for 1 hour at room temperature. Membrane was washed in PBS containing 0.1% Tween-20 (PBS-T) and probed with Drp1-specific mouse monoclonal antibody (1.0 µg/mL, clone 4E11B11, Cell Signalling) at 4°C overnight. Following successive washes with PBS-T, the membrane was probed with secondary antibody (IRDye 800CW Goat anti-mouse (0.05 µg/mL; LiCOR Biosciences) at room temperature for 1 hour. After washing, membrane was scanned at 800 nm wavelength using the Odyssey CLx scanner (LiCOR Biosciences), and reprobed with α-tubulin monoclonal antibody (0.36 µg/mL, clone DM1A; Cell Signalling) for 60 min at room temperature. Protein band intensity was determined by densitometry using Image Studio software (v.6).

### Mitochondrial morphology analysis

Mitochondrial morphology was assessed using Hsp60 immunostaining on MEFs cultured in 8-well chamber slides (4000 cells/well, Thermo Fisher Scientific) precoated with 0.1% (w/v) gelatin [33]. Fixed MEFs were stained with Anti-Hsp60 (2 µg/mL, rabbit polyclonal, ab46798, Abcam, lot#1071079-1) overnight at 4°C followed by Alexa Fluor-488 goat anti-rabbit secondary antibody (10 μg/mL, Invitrogen) and DAPI (1 μg/mL) for 1 hour at room temperature. Coverslips were mounted using fluorescence mounting medium (Dako). Images were acquired using an Olympus BX61 fluorescence microscope at 600x magnification. For each replicate, images were captured from 20 random fields of view, with a minimum of 200 cells analysed. Mitochondrial morphology was classified as either predominantly elongated (network) or fragmented (tubular), based on whether more than 50% of the mitochondria in a given cell exhibited that morphology. For each condition, the combined percentage of cells classified as having elongated or fragmented mitochondria totals 100%.

### Recombinant His_6_-Drp1 Expression and Purification

cDNAs encoding human Drp1 isoform 1 (UniProt ID: O00429-1), isoform 2 (UniProt ID: O00429-3), isoform 3 (UniProt ID: O00429-4), isoform 4 (UniProt ID: O00429-2), isoform 5 (UniProt ID: O00429-6), isoform 6 (UniProt ID: O00429-8), and isoform 8 (UniProt ID: O00429-9) were synthesised by General Biosystems with N-terminal His_6_-tags and cloned into the pET-21b(+) vector using *NdeI/NotI* restriction sites. All constructs were sequence-verified and transformed into *Escherichia coli* Rosetta (DE3) cells (Novagen, Merck Millipore). For recombinant protein expression, transformed cells were cultured in 2 L of 2× yeast extract–tryptone medium supplemented with 100 µg/mL ampicillin at 37°C, shaking at 120 rpm, until reaching an optical density of *A*_600_□∼1.0. Protein expression was induced with 0.5 mM β-D-1-thiogalactopyranoside, followed by incubation at 16°C for ∼16 hours. Cells were harvested by centrifugation at 3000 rpm at 4°C for 20 minutes.

Cell pellets were resuspended in ice-cold lysis buffer. For all isoforms except isoform 3, the lysis buffer contained 50 mM HEPES (pH 7.55), 400 mM NaCl, 5 mM MgCl_2_, 40 mM imidazole, 2.5 mM β-mercaptoethanol (BME), 10 µM leupeptin, 100 µM (4-(2-aminoethyl)benzenesulfonyl fluoride) (AEBSF), and 1 mM benzamidine hydrochloride. For isoform 3, the buffer contained 50 mM HEPES (pH 7.8), 400 mM NaCl, 5 mM MgCl_2_, 40 mM imidazole, 10 µM leupeptin, 100 µM AEBSF, and 1 mM benzamidine hydrochloride. Cells were lysed with a pre-chilled EmulsiFlex-C5 homogenizer (Avestin, Ottawa, Canada) and clarified by centrifugation at 16,500 rpm at 4°C for 30 minutes.

Supernatants were loaded onto Nickel Sepharose Fast Flow 6 resin (GE Healthcare, Buckinghamshire, UK) for affinity purification. For all isoforms except isoform 3, proteins were eluted in 50 mM HEPES (pH 7.55), 400 mM NaCl, 5 mM MgCl_2_, 400 mM imidazole, and 2.5 mM BME. For isoform 3, elution buffer contained 50 mM HEPES (pH 7.8), 400 mM NaCl, 5 mM MgCl_2_, 500 mM imidazole, and freshly added 5 mM BME. Eluted protein were subjected to size-exclusion chromatography on a HiLoad 16/600 Superdex 200 pg column (GE Healthcare) equilibrated in storage buffer. For all isoforms except isoform 3, storage buffer contained 20 mM HEPES (pH 7.5), 300 mM NaCl, 2.5 mM MgCl_2_, and 2.5 mM tris(2-carboxyethyl)phosphine (TCEP). For isoform 3, storage buffer contained 20 mM HEPES (pH 7.5), 300 mM NaCl, 2,5 mM MgCl_2_, and freshly added 5 mM BME. Purified Drp1 proteins were aliquoted, flash-frozen in liquid nitrogen, and stored at -80°C.

### Mass photometry

Mass photometry measurements of Drp1 were performed on the Two MP system (Refyn, MA, USA). The instrument was calibrated using bovine serum albumin and thyroglobulin. Measurements were performed in a total volume of 40 μL per well. For each experiment, 10 μL of MP buffer (20 mM HEPES (pH 7.5), 300 mM NaCl, 2.5 mM MgCl_2_, and 2.5 mM tris(2-carboxyethyl)phosphine (TCEP)) was added to clean wells to calibrate the focal height. Freshly thawed Drp1 protein was diluted in MP buffer, and 10 μL of the protein solution was added to the wells to achieve a final concentration of 200 nM, with and without 2 mM GMPPCP. Each measurement was recorded for 120 seconds, and at least three independent replicates were performed for each Drp1 isoform. Molecular masses of detected species between 65 kDa and 1200 kDa were determined using the calibration curve.

### Surface Plasmon Resonance (SPR)

All SPR experiments were performed using a Biacore T200 instrument (GE Healthcare Life Sciences, IL, USA) at 25 °C in HBS-EP+ buffer (20 mM HEPES, pH 7.4, 150 mM NaCl, 50 μM EDTA, and 0.005% Tween-20). Drp1 protein was tethered to a NiHC1500 SPR sensor chip (Xantec) via His-tag capture in SPR buffer. Incremental injections of the protein were performed to achieve a surface density of approximately 8000 RU (flow rate 10 μL/minute), corresponding to a calculated maximum analyte binding capacity (Rmax) of ∼45 RU. The immobilized protein surface was allowed to stabilise for at least 1 hour in HBS-EP+ buffer prior to analyte injection. GDP (Merck) was dissolved in HBS-EP+ buffer to prepare a 100 mM stock solution and subsequently diluted to desire cncentrations. The compound was injected across the chip surface at 30 μL/minute for 60 seconds, followed by 400 seconds dissociation phase. A threefold serial dilution series was used, starting from a maximum concentration of 120 μM (10-point dilution series). Data were analysed using the Biacore T200 Evaluation Software version 1.

### Drp1 GTPase Activity Assay

Malachite green reagent was prepared by mixing 0.0812% (w/v) malachite green (M-9636, Sigma-Aldrich) in Milli-Q water, 2.32% (w/v) polyvinyl alcohol (P8136-250, Sigma-Aldrich) in Milli-Q water (dissolved with heating), 5.72% (w/v) ammonium molybdate (A7302, Sigma-Aldrich) in 6 M HCl, and Milli-Q water at a volume ratio of 2:1:1:2. The mixture was incubated at room temperature in the dark for approximately 2 hours before use. Recombinant human Drp1 (0.5 µM for all isoforms, except for isoform 2, which was used at 0.25 µM) was incubated with 0.5 mM GTP (R0461, Thermo Scientific) in assay buffer containing 50 mM HEPES (pH 7.5), 2 mM MgCl_2_, and 1 mM dithiothreitol (DTT) at 37°C in the dark for 60 minutes, in a final volume of 50 µL in a 96-well microtiter plate. The concentration for each isoform was empirically determined to remain within the assay’s linear range. Reactions were terminated by addition 12.5 µL of 0.5 M EDTA (pH 8.0), followed by 50 μL malachite green reagent. Subsequently, 5 µL of 34% sodium citrate was added to each well to prevent nonenzymatic hydrolysis of ATP from contributing to background signal [34]. Plates were incubated at room temperature in the dark for 30 minutes. Inorganic phosphate release from GTP hydrolysis was quantified by measuring absorbance at 650 nm using a CLARIOstar Plus plate reader (BMG Labtech). Negative controls included reactions lacking either Drp1 or GTP. All assays were performed in triplicate, and each experiment was repeated independently at least three times.

### Statistics

All values are expressed as mean ± standard error of the mean (SEM). Significance of the differences was evaluated using one-way ANOVA followed by multiple comparison post-hoc analysis where appropriate. *p* ≤ 0.05 was considered statistically significant.

## Results

### Capture of full-length DNM1L/Dnm1l transcripts in human and mouse tissues

To investigate transcript diversity of Drp1, we performed high-specificity long-read sequencing using a tailored cDNA synthesis and amplification workflow optimized for full-length isoform resolution (**Figure 3A**). Starting from 300 ng of high-integrity total RNA (RIN >8) per sample, reverse transcription was carried out with Maxima H Minus Reverse Transcriptase using a custom oligo(dT) primer and a template-switch oligo containing a Unique Molecular Identifier (UMI), enabling precise molecule tracking and error correction. A key specificity-enhancing step involved targeted amplification of Drp1 transcripts using biotinylated primers (HEx21-R1 for human or MEx21-R1 for mouse) designed to anneal at exon 21 before the stop codon, a region conserved across all known isoforms. This targeted enrichment strategy, coupled with the use of high-fidelity KAPA HiFi polymerase and template-switch-based amplification, enabled selective capture of full-length Drp1 isoforms. Biotin-tagged amplicons were isolated via streptavidin-coated magnetic beads and re-amplified using the same exon-specific primers to ensure robustness and consistency. Purified libraries were sequenced using PromethION platform from Oxford Nanopore Technologies with native barcoding, and base calling was conducted in super accuracy mode, providing high-confidence long-read data. This targeted, UMI-guided, full-length approach enabled deep isoform-resolved analysis of Drp1, capturing transcript variants with high specificity and minimizing amplification bias.

Comprehensive data analysis confirmed the high quality and reproducibility of our targeted long-read sequencing pipeline. Quality control analyses were performed to confirm the robustness and lack of bias in the amplicon capture and sequencing workflow. The distribution of read lengths across samples and groups showed consistent profiles, with modal lengths corresponding to full-length Drp1 transcript isoforms, indicating successful amplification and capture of complete intact cDNA (**Figure 3B, Supplementary Figure 1**). UMI saturation analysis demonstrated near-complete recovery of unique transcript molecules within each sample, indicating that sequencing depth was sufficient and that PCR duplicates were effectively accounted for by UMI collapsing (**Supplementary Figure 2A**). Using human left ventricle samples as a reference, we performed correlation analysis between Drp1 transcript expression levels in our targeted amplicon dataset and public GTEx bulk long read RNA-seq data. The analysis revealed strong concordance (Pearson’s R = 0.89), supporting that isoform proportions are preserved in our targeted sequencing approach (**Figure 3C**). Furthermore, several lowly expressed transcripts were only detected using our approach, demonstrating the comprehensive capture of all isoforms of Drp1. Transcript coverage profiling from the 5’ to 3’ ends confirmed uniform representation across the transcript length with no positional bias, and a clear drop of coverage corresponded to the position of primer location and start of 5’ UTR. This underscores the effectiveness of template-switching combined with oligo(dT)-primed reverse transcription (**Supplementary Figure 3A-B**). Similar uniform representation was observed using exonic coverage, except exon 3, 16 and 17, consistent with high-efficiency capture of full-length isoforms (**Supplementary Figure 3C-D**). These depleted exons were consistent with the visual inspection of Drp1 in Integrated Genomics Viewer (IGV) in representative samples, which also illustrate clean mapping of reads, absence of spurious alignments, and accurate capture of exon-exon junctions (**Supplementary Figure 4**). Expression of three selected Drp1 isoforms showed strong correlation between human left ventricle and mouse heart tissues, supporting both technical reproducibility and biological conservation across species (**Figure 3D**).

### Transcript variant diversity of DNM1L in human cardiomyocytes and heart tissues

Our long-read RNA sequencing dataset demonstrates robust capability to capture full-length *DNM1L* transcripts in human iPSC-derived cardiomyocytes (iCM-CERA and iCM-CL2) and human left ventricle, enabling accurate isoform-level analysis. The on-target rate for the *DNM1L* gene ranged from 8.8% to 36.7% across samples (**Figure 4A**). There were 28,821-133,361 unique UMIs for *DNM1L t*ranscript in each sample (**Figure 4B**). Principle Component Analysis (PCA) based on log_2_-transformed Transcript unique UMI counts Per Million (log_2_TPM) for *DNM1L* transcripts showed expected sample clustering, with iPSC-derived cardiomyocytes separating slightly from primary left ventricle samples (**Figure 4C**). This suggests that Drp1 isoform diversity is associated with cardiac tissue identity and developmental state.

**Figure 4.**
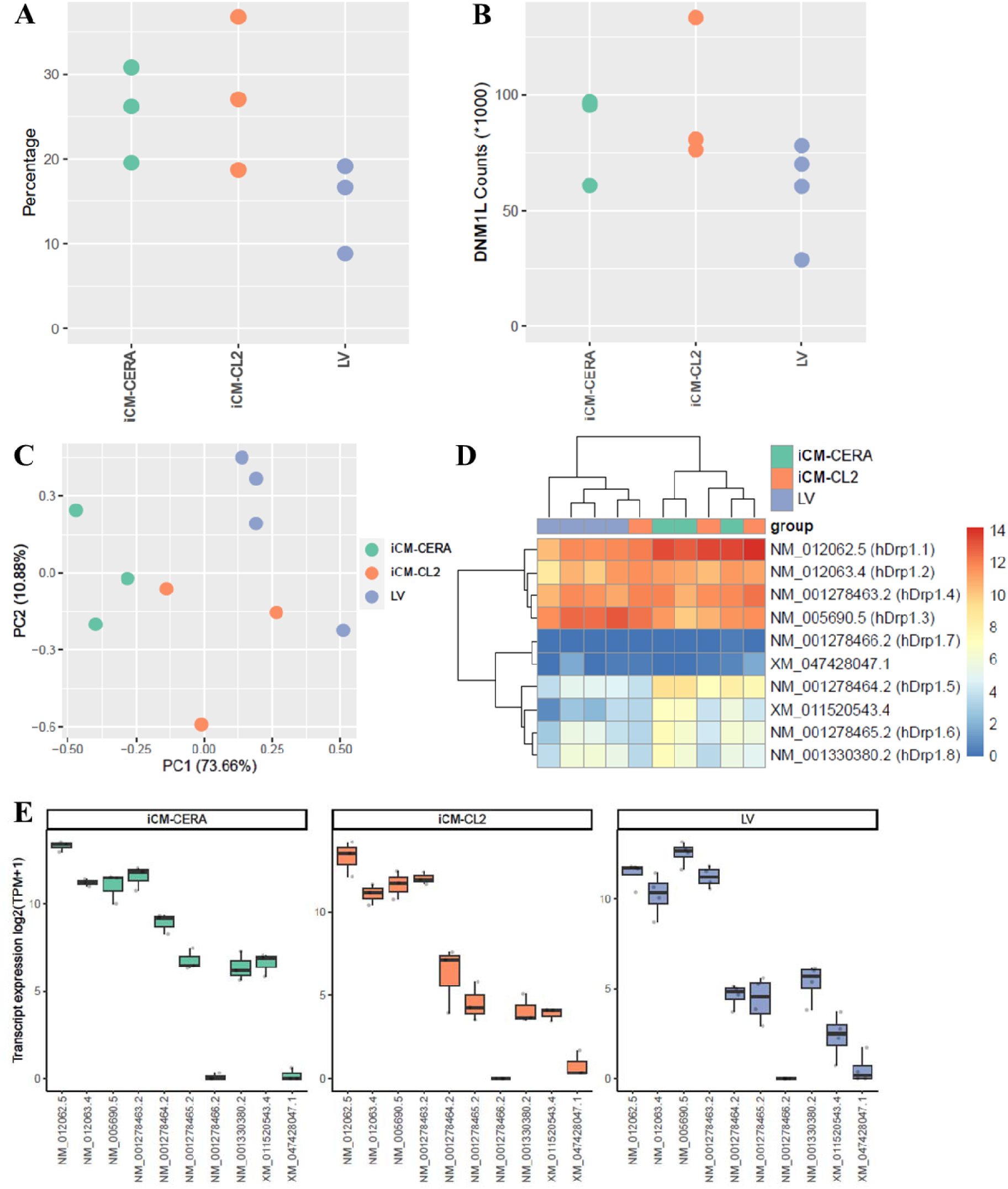
Expression of *DNM1L* transcript variants in human cardiomyocytes and heart tissue. **(A)** On-target rate for the *DNM1L* gene. **(B)** Total unique UMI counts for all *DNM1L* transcript. **(C)** PCA plot of iCM-CERA, iCM-CL2, and primary left ventricle samples based on *DNM1L* transcripts using log_2_(TPM+1). **(D)** Heatmap of *DNM1L* transcript expression [log_2_(TPM + 1)]. **(E)** Boxplots showing *DNM1L* transcript expression [log_2_(TPM+1)] in iCM-CERA, iCM-CL2, and primary left ventricle samples.

The dataset comprehensively recovered the full set of annotated Drp1 (*DNM1L*) transcripts listed in the NCBI record. In both human iPSC-derived cardiomyocytes and primary left ventricle samples, transcript variant abundance patterns were broadly similar. Isoforms 1, 2, 3, and 4, all lacking exon 3, were the most abundantly expressed across both cell types, followed by moderate expression of isoforms 5, 6, and 8. Isoform 7, missing exon 5 to 8, was consistently detected at low levels. Among the two XM transcripts identified, XM_011520543.4 displayed moderate expression, whereas XM_047428047.1 was expressed at low levels, comparable to isoform 7 (**Figure 4D-E**). Together, these results confirm the robust capture of the full spectrum of DNM1L transcript variants in both cardiomyocytes and primary heart tissue.

The functional relevance of the observed expression patterns was assessed by measuring the oligomerization state, GTP-binding affinity, and GTPase activity of recombinant Drp1 isoforms. Drp1 mediates mitochondrial fission through oligomerisation on the outer mitochondrial membrane. At a concentration of 200 nM, Drp1 isoforms 1, 2, 3, 4, 5, 6, and 8 exhibited comparable oligomerisation behaviour, existing predominately as dimers both in solution (**Figure 5A**) and in the presence of the non-hydrolysable GTP analogue GMPPCP (data not shown). All isoforms bound GDP with low micromolar affinity (**Figure 5B**). Among them, isoform 3 showed noticeably weaker GDP binding, as indicated by a higher KD value relative to the other isoforms (**Figure 5B**).

**Figure 5.**
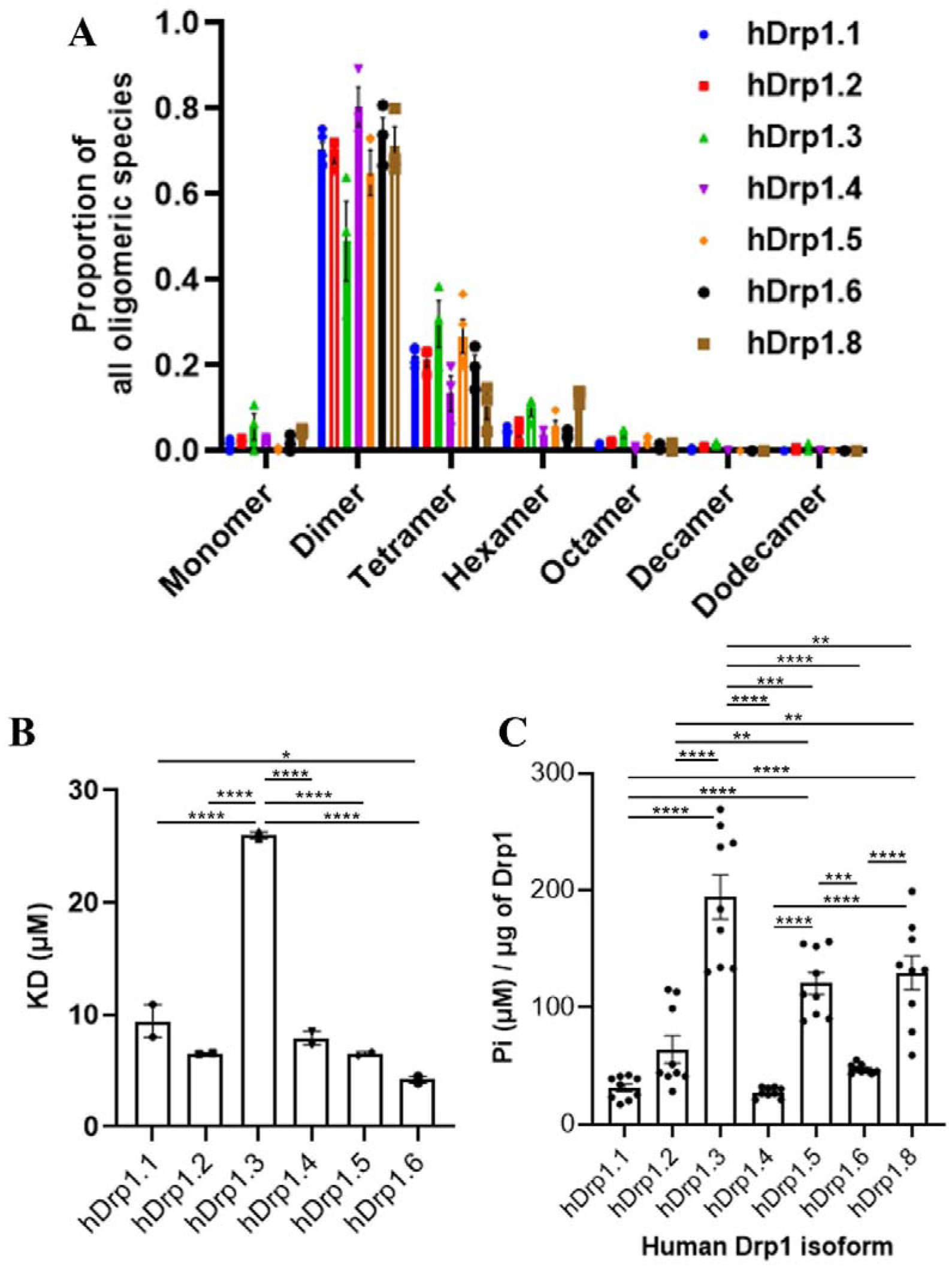
Biochemical analysis of recombinant human Drp1 isoform proteins. Recombinant Drp1 isoforms were expressed, purified, and analysed for their oligomerisation state **(A)**, GDP-binding affinity **(B)**, and GTPase enzymatic activity **(C)**. **(A)** Oligomerization analysis showed that all isoforms were predominantly dimeric at a concentration of 200 nM. Measurements were performed in triplicate and independently repeated at least three times. **(B)** SPR analysis of GDP binding revealed that isoform 3 exhibited weaker binding affinity compared to the other isoforms. Assay were performed in duplicate and independently repeated twice. **(C)** GTPase activity was quantified by measuring the rate of inorganic phosphate release over time in the presence of GTP. All reactions were conducted under identical buffer and temperature conditions. Assays were performed in triplicate and independently repeated three times. Data are presented as mean ± SEM. **p < 0.01, ***p < 0.001, ****p < 0.0001 by one-way ANOVA with Bonferroni post hoc test.

Analysis of enzymatic activity revealed that isoforms 3, 5, and 8 exhibited the highest enzymatic activity, whereas isoforms 1, 2, 4, and 6 showed comparatively lower activity (**Figure 5C**). Thes findings indicate that transcript abundance does not necessarily correlate with biochemical activity of Drp1 isoforms. The findings highlight the importance of integrating both expression data and biochemical characterisation to understand isoform-specific contributions to Drp1 function in the cardiomyocytes and heart tissues.

### Long-read RNA sequencing of Dnm1l transcript variants in primary mouse tissues

Long-read RNA sequencing enabled comprehensive detection of *Dnm1l* (DRP1) isoforms across multiple primary mouse tissues, including brain, heart, kidney, lung, skeletal muscle, and spleen. The on-target rate for the *Dnm1l* gene ranged from 15.8% to 51.8% across samples (**Figure 6A**). There were 59,584 to 178,684 unique UMIs for *Dnm1l* transcripts in each sample, providing robust coverage. The dataset recovered the complete set of annotated *Dnm1l* transcripts listed in the NCBI record, encompassing 22 NM (protein-coding), 14 NR (non-coding RNA), and one XM (predicted protein-coding) transcript (**Figure 6**), underscoring the sensitivity and completeness of long-read sequencing in capturing both coding and non-coding variants across the locus. These results confirm effective detection of all known isoforms, ensuring sufficient depth and diversity for isoform-resolved analyses.

**Figure 6.**
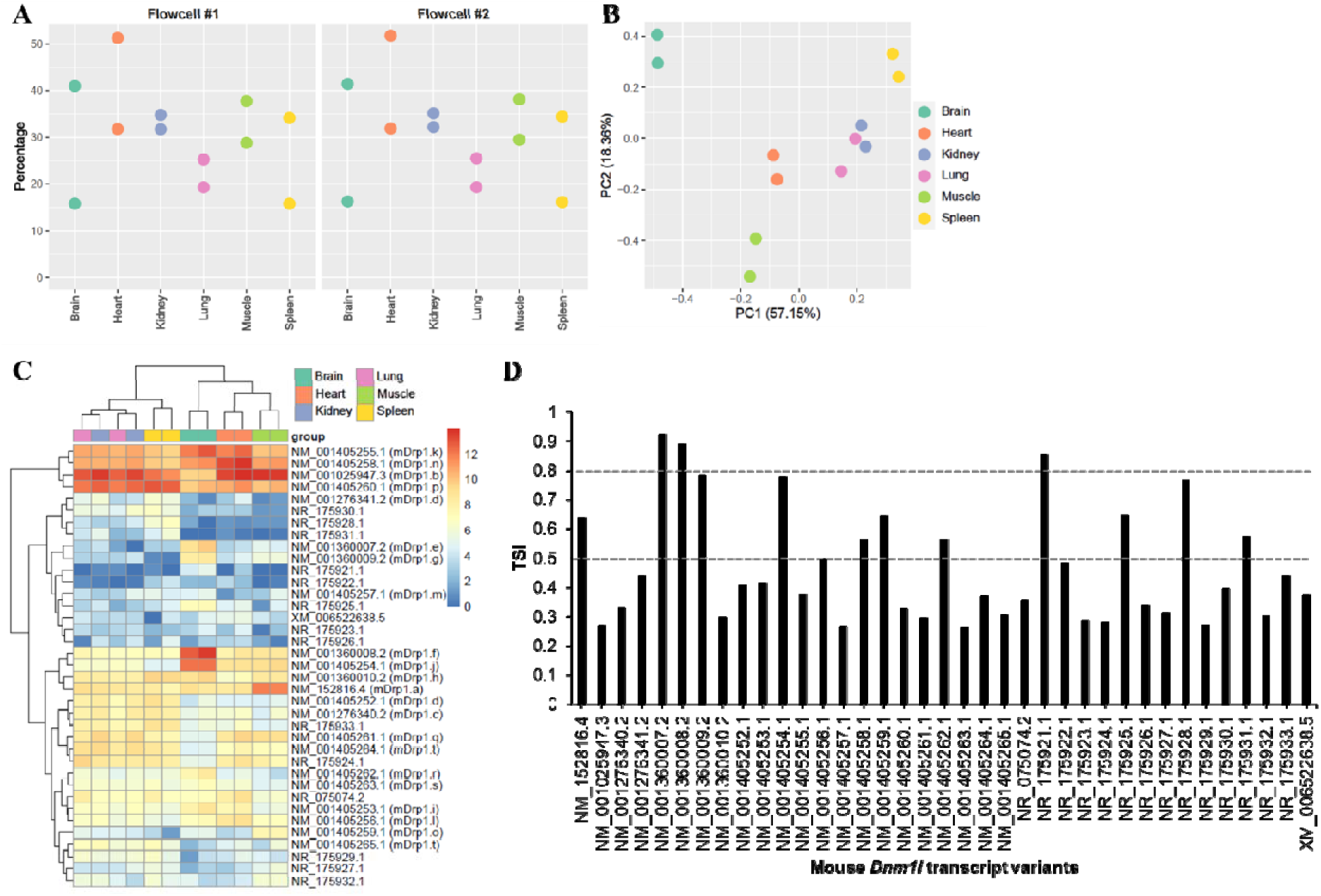
Expression of *Dnm1l* transcript variants in mouse tissues. **(A)** On-target rate for the Dnm1l gene. **(B)** PCA plot of mouse brain, heart, kidney, lung, skeletal muscle and spleen samples. **(C)** Heatmap of *Dnm1l* transcript expression [log_2_(TPM + 1)]. **(D)** Tissue specificity index (TSI) of all Dnm1l transcripts across mouse tissues.

PCA revealed distinct tissue-specific clustering of *Dnm1l* transcripts (**Figure 6B**). Brain and spleen occupied separate regions in the PCA space, while organs rich in epithelial tissues, such as lung and kidney, clustered together. Heart and skeletal muscle were positioned in close proximity, reflecting similarities in their *Dnm1l* transcript profiles. Overall, *Dnm1l* transcript expression contributed substantially to the principal components, suggesting that isoform diversity correlates with tissue identity.

A more detailed analysis of *Dnm1l* isoform expression revealed a spectrum of transcriptional patterns, ranging from broadly expressed to highly tissue-specific isoforms (**Figure 6C**). Hierarchical clustering of tissues and isoforms identified groups of co-expressed transcripts, consistent with potential functional specialisation of specific Drp1 isoforms. Isoforms e (NM_001360007.2), f (NM_001360008.2), g (NM_001360009.2) and j (NM_001405254.1) were predominantly expressed in brain, suggesting a specialised role in neuronal function. In contrast, isoforms l (NM_001405256.1) and n (NM_001405258.1) were selectively enriched in heart, while isoforms a (NM_152816.4) and o (NM_001405259.1) were relatively enriched in skeletal muscle. Boxplots of individual transcript expression revealed both within- and between-tissue variability, confirming the existence of dominant isoforms with broad expression alongside minor isoforms with restricted, tissue-specific expression (**Supplementary Figure 5**). Tissue Specificity indices (TSI) analysis quantitatively supported these findings, with values near 1 indicating highly tissue-specific isoforms and values near 0 reflecting broadly expressed transcripts (**Figure 6D**).

Together, these analyses demonstrate that long-read RNA sequencing not only captures the full complement of canonical *Dnm1l* transcripts but also resolves isoform-specific expression dynamics across tissues. The observed tissue-enriched expression of select isoforms suggests that Drp1 isoform diversity is tightly linked to tissue identity and may underlie specialized regulation of mitochondrial dynamics in distinct cellular contexts.

### Differential effects of Drp1 isoforms on mitochondrial fission

To investigate the specific contributions of Drp1 isoforms to the regulation of mitochondrial morphology, a Drp1 knockout (KO) mouse embryonic fibroblast (MEF) model was used (**Figure 7**). Distinct mouse Drp1 splice variants were individually and stably expressed in theses cells to evaluate their ability to rescue mitochondrial fission defects. The isoforms tested included: **isoform b**, which lacks exons 3, 16, and 17 and contains an incomplete exon 2, and is broadly expressed across tissues; **isoform d**, which lacks exons 1-4 and 16; **isoform e**, the full-length canonical form that is highly expressed in brain; and **isoform o**, which lacks exons 16 and 17, and is enriched in skeletal muscle. Sequence homology analysis using BLASTp revealed that mouse Drp1 isoforms b, d, e, and o exhibited the highest similarity with human Drp1 isoforms 3, 2, 5, and 8, respectively (**Table 3**). Western blotting revealed similar stable protein expression of mouse isoforms b, e and o in KO MEFs, with expression of isoform d detected at noticeably lower levels (**Figure 7A**).

**Figure 7.**
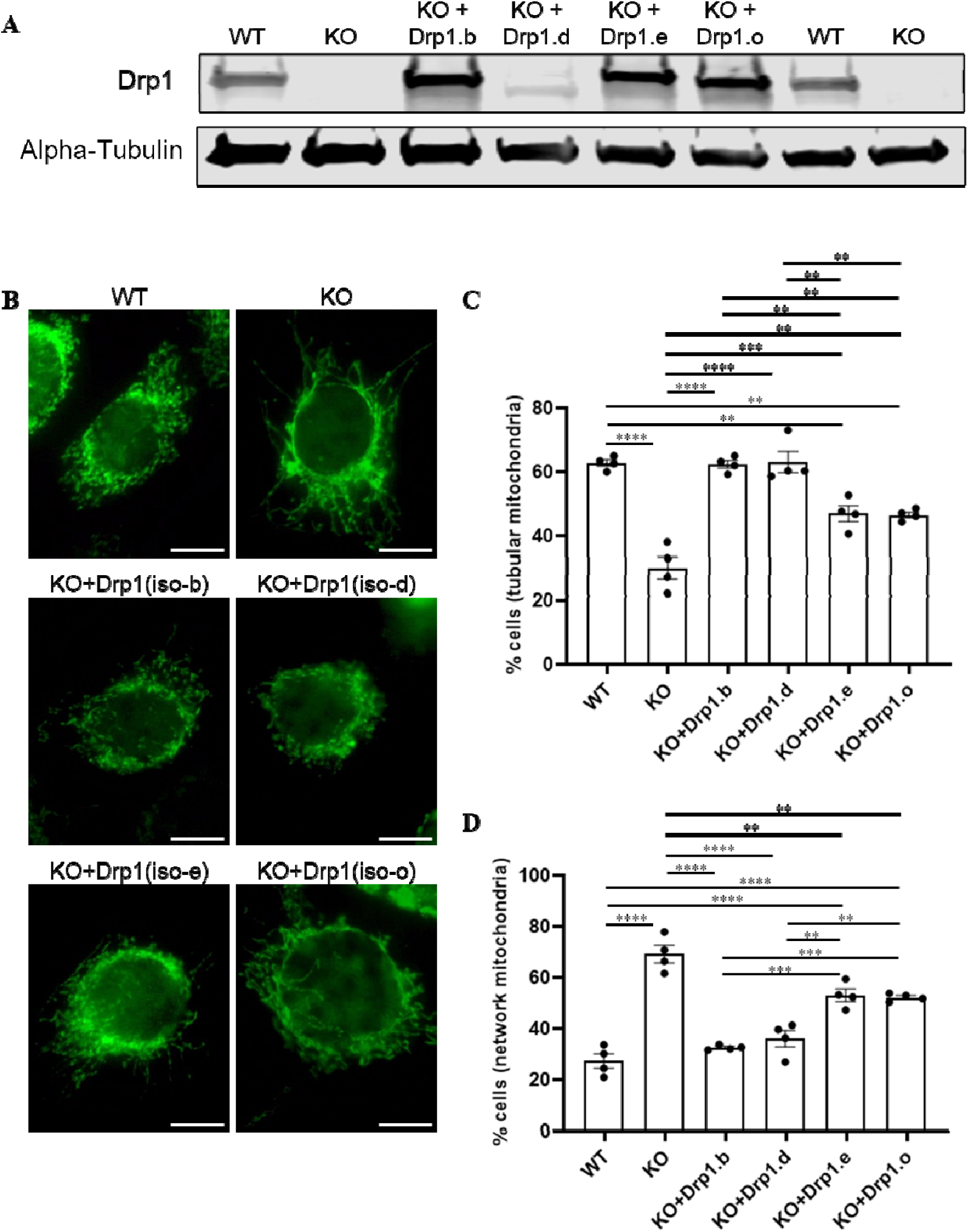
Mitochondrial morphology in MEFs. **(A)** Protein expression of Drp1 in KO MEFs. **(B)** Representative fluorescent images of mitochondrial morphology at 600x magnification in MEFs stained with Hsp60 antibody. Scale bar = 10 μm. **(C-D)** The percentage of cells with **(C)** tubular or **(D)** network mitochondria in MEFs (wild type, Drp1-KO, Drp1-KO re-expressing Drp1 isoform-b, isoform-d, isoform-e or isoform-o). N=4. Data are expressed as mean ± SEM. **p < 0.01, ***p < 0.001, ****p < 0.0001 by one-way ANOVA with Bonferroni post hoc test

**Table 3:** Comparative sequence similarity of Drp1 isoforms in mouse and human using BLASTp algorithm.

| Mouse Drp1 isoforms |  | Max Score (Percent Identity/Percent Query Cover) |  |  |  |  |  |  |  |
| --- | --- | --- | --- | --- | --- | --- | --- | --- | --- |
| Isoform | Transcript Protein Accession | Human isoform 1 | Human isoform 2 | Human isoform 3 | Human isoform 4 | Human isoform 5 | Human isoform 6 | Human isoform 7 | Human isoform 8 |
| a | NP_690029.2 | 1391 (92.52/100) | 1400 (95.57/100) | 1408 (97.05/100) | 1395 (93.90/100) | 1431 (94.26/100) | 1434 (95.66/100) | 912 (91.84/91) | 1446 (98.88/100) |
| b | NP_001021118.1 | 1410 (94.16/100) | 1410 (97.32/100) | 1417 (98.86/100) | 1406 (95.59/100) | 1391 (92.52/100) | 1395 (93.90/100) | 912 (91.84/91) | 1408 (97.05/100) |
| c | NP_001263269.1 | 1418 (94.47/100) | 1431 (97.91/100) | 1402 (96.51/100) | 1398 (94.53/100) | 1415 (93.47/100) | 1394 (93.50/100) | 936 (93.47/91) | 1399 (95.44/100) |
| d | NP_001392181.1 | 1230 (94.82/100) | 1243 (98.85/86) | 1214 (97.22/85) | 1210 (94.89/86) | 1230 (94.82/85) | 1209 (94.89/84) | 935 (93.47/91) | 1214 (97.22/84) |
| e | NP_001346936.1 | 1478 (96.16/100) | 1414 (92.85/100) | 1389 (91.52/100) | 1449 (94.83/100) | 1508 (97.75/100) | 1479 (96.42/100) | 998 (98.57/91) | 1419 (93.11/100) |
| f | NP_001346937.1 | 1481 (96.96/100) | 1416 (93.59/100) | 1390 (92.26/100) | 1449 (94.83/100) | 1518 (98.66/100) | 1488 (97.33/100) | 999 (98.57/91) | 1430 (93.99/100) |
| g | NP_001346938.1 | 1448 (94.83/100) | 1394 (93.01/100) | 1392 (92.88/100) | 1454 (96.24/100) | 1479 (96.42/100) | 1488 (97.85/100) | 970 (96.53/91) | 1424 (94.49/100) |
| h | NP_001346939.1 | 1416 (93.59/100) | 1429 (96.96/100) | 1400 (95.57/100) | 1394 (93.63/100) | 1455 (95.33/100) | 1434 (95.39/100) | 936 (93.47/91) | 1438 (97.37/100) |
| i | NP_001392182.1 | 1482 (97.84/100) | 1418 (94.47/100) | 1394 (93.13/100) | 1452 (96.50/100) | 1479 (96.80/100) | 1450 (95.47/100) | 998 (98.57/91) | 1390 (92.13/100) |
| j | NP_001392183.1 | 1481 (96.93/100) | 1396 (93.77/100) | 1394 (93.63/100) | 1459 (97.02/100) | 1488 (97.33/100) | 1498 (98.78/100) | 970 (96.53/91) | 1434 (95.39/100) |
| k | NP_001392184.1 | 1488 (98.64/100) | 1425 (95.24/100) | 1401 (93.89/100) | 1459 (97.28/100) | 1480 (96.93/100) | 1449 (95.59/100) | 999 (98.57/91) | 1390 (92.26/100) |
| l | NP_001392185.1 | 1453 (96.50/100) | 1401 (94.66/100) | 1398 (94.53/100) | 1461 (97.95/100) | 1451 (95.47/100) | 1456 (96.89/100) | 969 (96.53/91) | 1394 (93.50/100) |
| m | NP_001392186.1 | 1396 (92.85/100) | 1427 (96.16/100) | 1397 (94.79/100) | 1375 (91.36/100) | 1445 (94.44/100) | 1423 (94.49/100) | 936 (93.47/91) | 1429 (96.43/100) |
| n | NP_001392187.1 | 1459 (97.28/100) | 1406 (95.45/100) | 1405 (95.31/100) | 1469 (98.76/100) | 1451 (95.59/100) | 1458 (97.02/100) | 970 (96.53/91) | 1394 (93.63/100) |
| o | NP_001392188.1 | 1389 (91.79/100) | 1398 (94.79/100) | 1405 (96.24/100) | 1377 (92.34/100) | 1422 (93.38/100) | 1425 (94.76/100) | 910 (91.84/91) | 1437 (97.91/100) |
| p | NP_001392189.1 | 1425 (95.24/100) | 1437 (98.73/100) | 1409 (97.32/100) | 1405 (95.31/100) | 1416 (93.59/100) | 1394 (93.63/100) | 936 (93.47/91) | 1400 (95.57/100) |
| q | NP_001392190.1 | 1395 (93.40/100) | 1402 (96.51/100) | 1410 (98.01/100) | 1398 (94.80/100) | 1391 (92.40/100) | 1394 (93.78/100) | 911 (91.84/91) | 1406 (96.91/100) |
| r | NP_001392191.1 | 1294 (98.74/86) | 1230 (94.82/86) | 1206 (93.25/85) | 1264 (97.17/86) | 1293 (98.74/85) | 1264 (97.17/84) | 998 (98.57/91) | 1205 (93.25/84) |
| s | NP_001392192.1 | 1265 (97.17/86) | 1211 (95.05/86) | 1210 (94.89/85) | 1273 (98.88/86) | 1264 (97.17/85) | 1273 (98.88/84) | 970 (96.53/91) | 1209 (94.89/84) |
| t | NP_001392193.1 | 1207 (93.56/86) | 1215 (97.22/86) | 1223 (99.00/85) | 1210 (95.21/86) | 1205 (93.56/85) | 1209 (95.21/84) | 911 (91.84/91) | 1222 (99.00/84) |

Mitochondrial morphology was assessed by quantifying the percentage of cells displaying predominantly tubular mitochondria or networked mitochondria, as a measure of fission activtiy. Compared to wild-type MEFs, Drp1-KO MEFs showed a significant reduction in mitochondrial fragmentation. Re-expression of isoform b or isoform d fully restored mitochondrial fragmentation, indicating robust functional capacity despite, in the case of isoform d, apparent low expression. In contrast, re-expression of isoform e and isoform o only partially rescued the fission defect (**Figure 7B-D**). These findings demonstrate that different Drp1 isoforms exert distinct effects on mitochondrial fission. Isoforms b and d, which both lack the A-insert within the GTPase domain (encoded by exon 3), were more effective in restoring fission than the full-length isoform e or the B-insert truncated isoform o, both of which retain the complete GTPase domain. Collectively, these data indicate that exons 2 and/or 3 may exert an inhibitory influence on Drp1-mediated mitochondrial fission.

## Discussion

Drp1, encoded by the *DNM1L* (dynamin-1-like) gene, is an evolutionarily conserved member of the dynamin family and a key regulator of mitochondrial fission. It is ubiquitously expressed, with particularly high levels in mitochondria-rich tissues such as the brain and heart. Dysregulation of Drp1 has been implicated in various pathophysiological conditions, including neurodegenerative diseases, cardiovascular disorders, and cancer [1]. Despite the established importance of Drp1 in mitochondrial dynamics and cell fate decisions, the diversity and functional significance of Drp1 isoforms remain incompletely understood. This knowledge gap is largely due to the limitations of conventional transcriptomic approaches. Mainstream short-read RNA sequencing, which involves RNA fragmentation prior to sequencing, is poorly suited to capturing full-length transcripts and accurately resolving transcript structures, particularly for variants involving widely spaced exons, such as the connections between exon 2–4 and exon 16–17 in *DNM1L*. Even advanced computational tools struggle to reconstruct complete isoform architectures from fragmented reads [35, 36], making it difficult to accurately determine the expression levels and specific functions of individual Drp1 isoforms under various pathophysiological conditions.

To overcome these limitations, the present study employs long-read sequencing technologies that preserve full-length RNA molecules, enabling high-resolution transcript-level analysis. Using targeted Nanopore native barcoded sequencing, we generated a comprehensive transcriptomic profile of *DNM1L* transcript variants in both human and mouse tissues. This approach allows us to accurately identify and quantify full-length *DNM1L* transcript variants, offering new insights into their potential roles in health and disease. Our analysis focused on alternative splicing events in exons 1–4, 16, and 17, which give rise to several coding variants annotated in the NCBI record [5, 6, 7, 14, 37, 38, 39]. Previous PCR-based studies reported tissue-specific expression of *DNM1L* transcript variants with alternative splicing of exons 16 and 17 in both healthy tissues and human brain tumours [6, 7]. By contrast, our long-read dataset captured isoforms spanning all 21 exons, revealing tissue-enriched rather than strictly tissue-specific expression of *DNM1L* transcript variants. For example, in mouse brain tissue, isoforms f, j, k, and n were expressed at higher levels than other isoforms. When expression was compared across tissue types, isoforms e, f, g, and j, each containing exon 3, were preferentially enriched in brain, suggesting a specialized neuronal role for exon 3-containing Drp1 variants. This expression pattern aligns with previous findings by Strack and colleagues, who reported predominant brain expression of exon 3-containing variants [5], and by Uo et al., who identified neuron-specific Drp1 variants incorporating exon 3 [14]. Interestingly, while Uo et al. suggested that exon 17-containing variants are absent in neurons [14], we observed substantial expression of isoforms e, f, k, and p, which contain exon 17, in mouse brain tissue. This discrepancy may reflect differences in sample composition where our dataset represents brain tissue containing diverse cell populations, whereas Uo and colleagues examined isolated neuronal cells.

In the heart, Drp1 isoforms l and n, which contain exon 16 but lack exons 3 and 17, are expressed at relatively higher levels compared with other tissue types in our dataset. This pattern contrasts with a previous report that described predominant expression of isoforms lacking exon 16 but containing exon 17 in the heart, kidney, and spleen [5]. Consistent with that study, however, we also observed that isoforms such as c and p, which lack exon 16 but include exon 17, were more abundant in the kidney and spleen. Notably, mouse *Dnm1l* transcript variants b (homologous to human isoform 3), k (homologous to isoforms 1 and 4), p (homologous to isoform 2), and n (homologous to isoform 4) were highly expressed in mouse heart tissue. This mirrors the strong expression of corresponding isoforms 1-4 in human heart tissue and iPSC-derived cardiomyocytes, suggesting that the regulation of these isoforms is conserved across species. Our findings reinforce the broader principle that Drp1 isoform diversity is functionally meaningful. While expression patterns show that isoforms are not restricted to a single tissue, the relative abundance and exon composition point to tissue-preferential functions. Such distribution suggests that isoform-specific structural differences adapt mitochondrial dynamics to the metabolic and structural requirements of different tissues.

Emerging evidence indicates that Drp1 isoforms exert distinct roles in apoptosis and mitochondrial morphology. Strack et al. demonstrated that MEFs lacking endogenous Drp1, as well as MEFs expressing Drp1-001 (lacking exons 3 and 16), were significantly less sensitive to staurosporine compared with MEFs expressing Drp1-000 (lacking exons 3, 16, and 17) or Drp1-011 (lacking exon 3). This reduced sensitivity was associated with a more fused mitochondrial network and a lower rate of apoptosis, suggesting that Drp1-001 (equivalent to hDrp1.2 and mDrp1.d) may promote cell survival by modulating mitochondrial dynamics [5]. Similarly, Uo et al. (2009) showed that only isoforms containing both exons 16 and 17 could interact with the anti-apoptotic protein Bcl-xL, underscoring isoform-specific protein-protein interactions in apoptotic regulation [14].

Our findings further indicate that alternative splicing of exon 3 modulates Drp1’s pro-fission activity. Isoforms lacking exon 3 (b and d) fully restored mitochondrial fragmentation in Drp1-null MEFs, whereas exon 3-containing isoforms (e and o) only partially rescued fission, suggesting that exon 3 inclusion exerts an inhibitory effect on Drp1-mediated fission. This aligns with earlier reports that exon 3 influences GTPase activity and protein interactions [8, 14], supporting the idea that exon 3-containing isoforms fine-tune, rather than maximize, Drp1 function. Their preferential enrichment in brain tissue supports the notion that these isoforms are tailored to the high metabolic demands and complex mitochondrial dynamics required for neuronal activity and plasticity. Although overexpression studies indicate that exon 3 does not determine Bcl-xL binding, SUMOylation, or intracellular localization [14], its contribution to GTPase modulation highlights the need for further functional characterization under physiological conditions.

In cancer, Drp1 isoform regulation is also critical. Javed et al. reported that high expression of a splice variant lacking exon 16 (Drp1(-/17)) is associated with poor prognosis in ovarian cancer patients [7]. Unlike the canonical exon 16/17-containing isoform, which localizes to mitochondrial fission puncta and promotes apoptosis, Drp1(-/17) mislocalises to microtubules [5, 7], leading to mitochondrial hyperfusion, enhanced oxidative metabolism, and resistance to pro-fission stimuli. These changes support tumour growth, survival, and chemoresistance, and manipulation of endogenous isoform ratios confirmed that Drp1(-/17) promotes proliferation and migration, whereas Drp1(16/17) limits tumour progression [7]. Interestingly, our *in vitro* assays did not fully align with these observations. Specifically, isoforms lacking exon 16 (hDrp1.3 and hDrp1.8) displayed significantly higher GTPase activity compared with exon 16-containing isoforms (hDrp1.1, hDrp1.4, and hDrp1.6). While these results do not contradict previous reports that exon 16-deficient isoforms mislocalise to microtubules and impair mitochondrial fission, they indicate that such isoforms possess elevated GTPase activity, which would theoretically be expected to facilitate, rather than suppress, mitochondrial fission. Thus, while clinical evidence points to a pro-tumorigenic role of exon 16-lacking variants, their precise mechanistic contribution to Drp1 enzymatic activity and mitochondrial dynamics remains to be clarified. Taken together, the relative abundance of exon 3-, exon 16-, and exon 16/17-containing variants emerges as a key determinant of mitochondrial morphology, apoptosis, and disease outcomes.

Our study demonstrates that Drp1 (Dnm1l) isoforms exhibit distinct GTPase activities and mitochondrial fission capacities, with expression patterns ranging from broadly expressed to highly tissue specific. By leveraging long-read RNA sequencing, we resolved full-length transcripts and captured a comprehensive isoform landscape, revealing both shared and tissue-restricted expression programs that reflect the specialized regulation of mitochondrial dynamics across cell types. These findings underscore the functional complexity of Drp1, showing that alternative splicing shapes isoform-specific roles in mitochondrial fission, apoptosis, metabolism, and disease. Rather than acting solely as a general mediator of mitochondrial division, Drp1 functions in a nuanced, isoform-dependent manner, with distinct contributions to cellular homeostasis and pathology. While the present study focused on annotated *Dnm1l* transcripts listed in the NCBI reference record, future investigations should leverage long-read sequencing technologies to uncover novel isoforms that are not yet captured in current annotations. Characterising these variants will provide a more comprehensive understanding of isoform-specific regulation, structural features, and interaction networks, which will be critical for elucidating their physiological roles and for evaluating the therapeutic potential of selectively targeting individual isoforms in neurodegeneration, cardiovascular disease, and cancer.

## Data Availability

The long-read Nanopore RNA sequencing data generated in this study have been deposited in the Gene Expression Omnibus under accession number GSE308731 (https://www.ncbi.nlm.nih.gov/geo/query/acc.cgi?acc=GSE308731). GTEx bulk long-read RNA sequencing data was downloaded from dbGAP under accession number phs000424.v9.

## Author contributions

SYL conceived the project, designed, and performed the experiments, and analysed and interpreted data. FY, JJ, NXYL, AMK, and LZ planned and performed experiments and analysed data. AAR, CL, JGL, SB, DAO, KL, and SL provided materials and methodologic input. NMD supervised the data analysis. FY, JH, NMD, JSO, and SYL provided interpretative review of study findings. All authors participated in manuscript preparation and approved the final version of the manuscript.

