## Supplementary materials for "Long-read sequencing-based atlas of tissue-specific expression of Drp1 transcript variants"

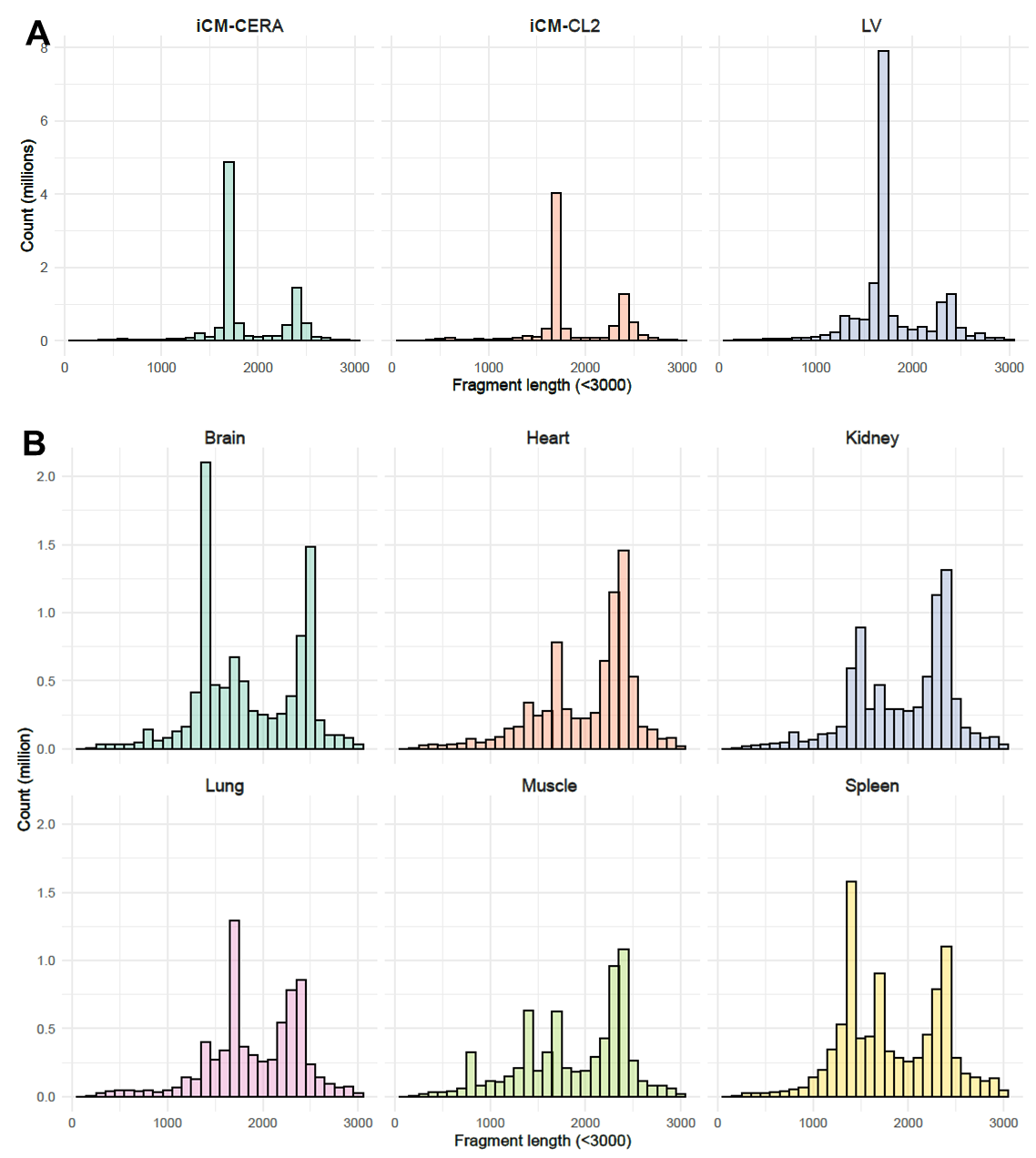

**Supplementary Figure 1.** Read length distributions across biological human **(A)** and mouse **(B)** samples.

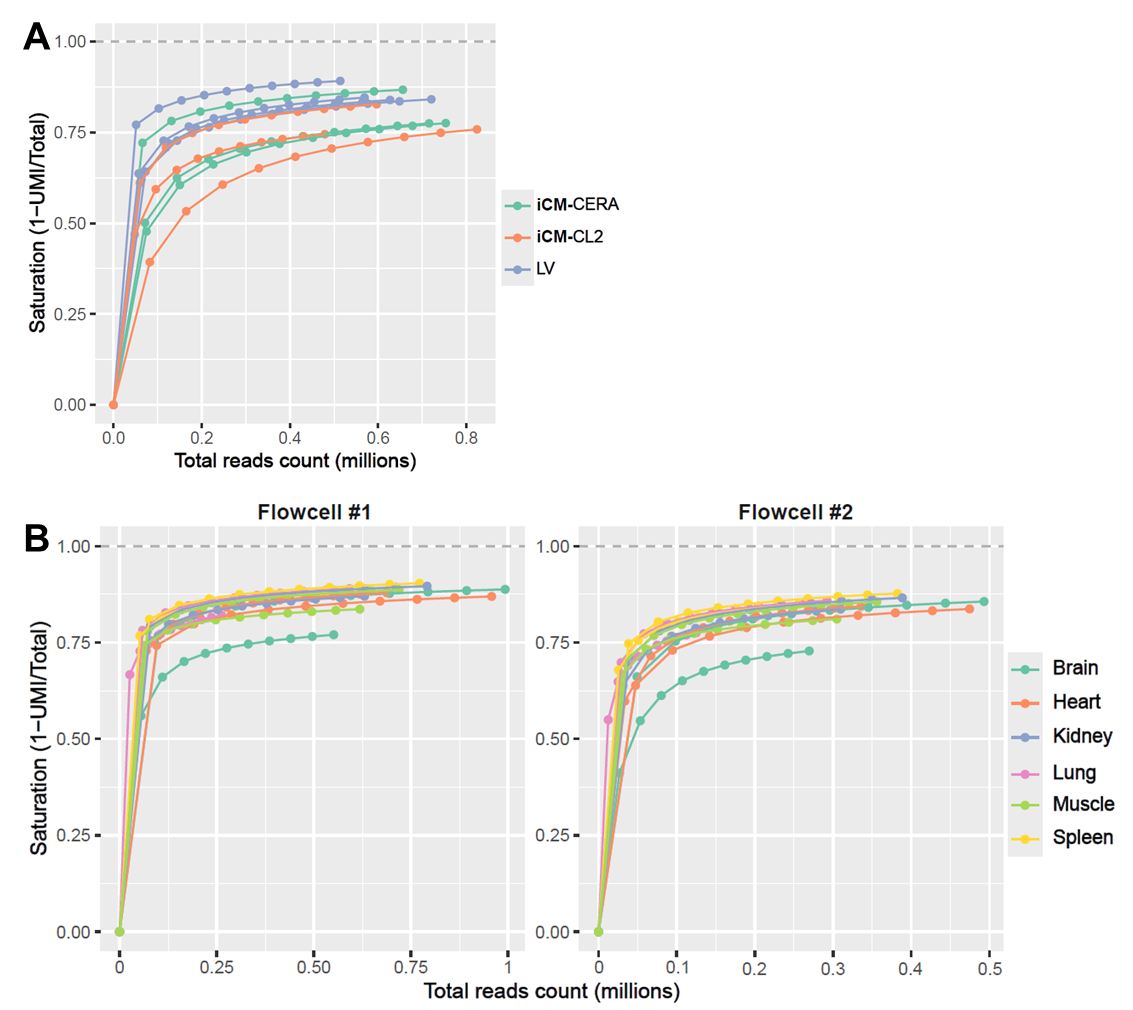

**Supplementary Figure 2.** Unique Molecular Identifier (UMI) saturation curve of human **(A)** and mouse **(B)** samples. For each sample, sequencing data was subsampled to defined percentages of the total mapped reads. UMI deduplication was then performed on each subsampled dataset to determine the number of unique UMI reads detected at each depth. Mouse data from two flowcells are shown separately.

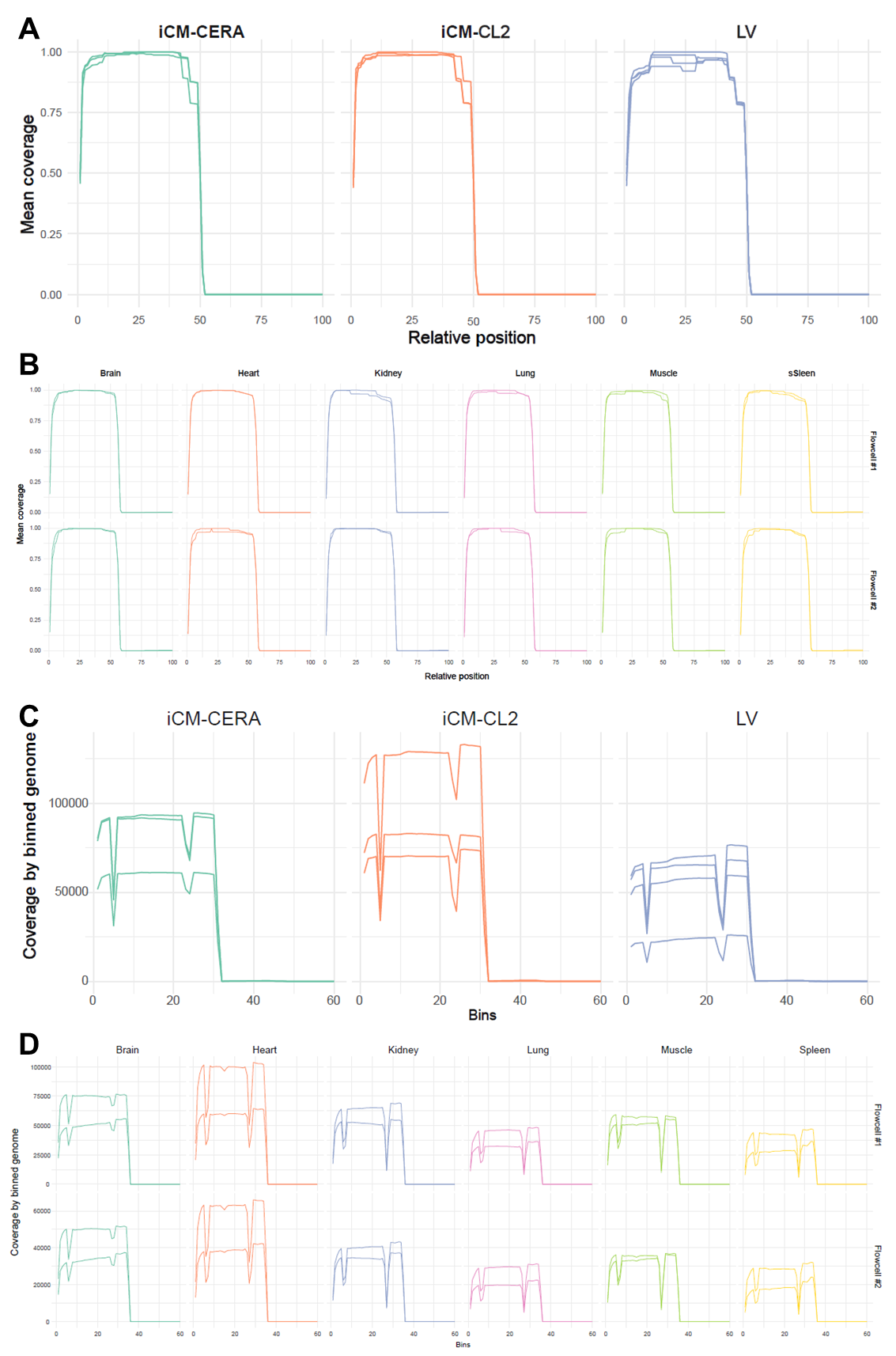

**Supplementary Figure 3. (A-B)** 5’–3’ transcript coverage across binned position in human **(A)** and mouse **(B)** samples. Mouse data from two flowcells are shown separately. For each transcript, coverage was calculated as the read depth within each bin normalized by the total reads aligned to that transcript. Coverage profiles of all *DNM1L* transcripts were averaged and plotted on the y-axis. The sharp drop in coverage corresponds to the location of the designed probes (~ 2200 bp from the 5’ end, near exon 21, representing ~50% of the transcript length). The downstream ~50% that is underrepresented corresponds primarily to the 3′ UTR. **(C-D)** Exonic coverage by binned position in human **(C)** and mouse **(D)** samples. Mouse data from two flowcells are shown separately. The coverage shown in y axis is the number of reads at each bin of concatenated human *DNM1L* 21 exon sequences. The drop of the line aligns with the location of exon 3 and 16, 17.

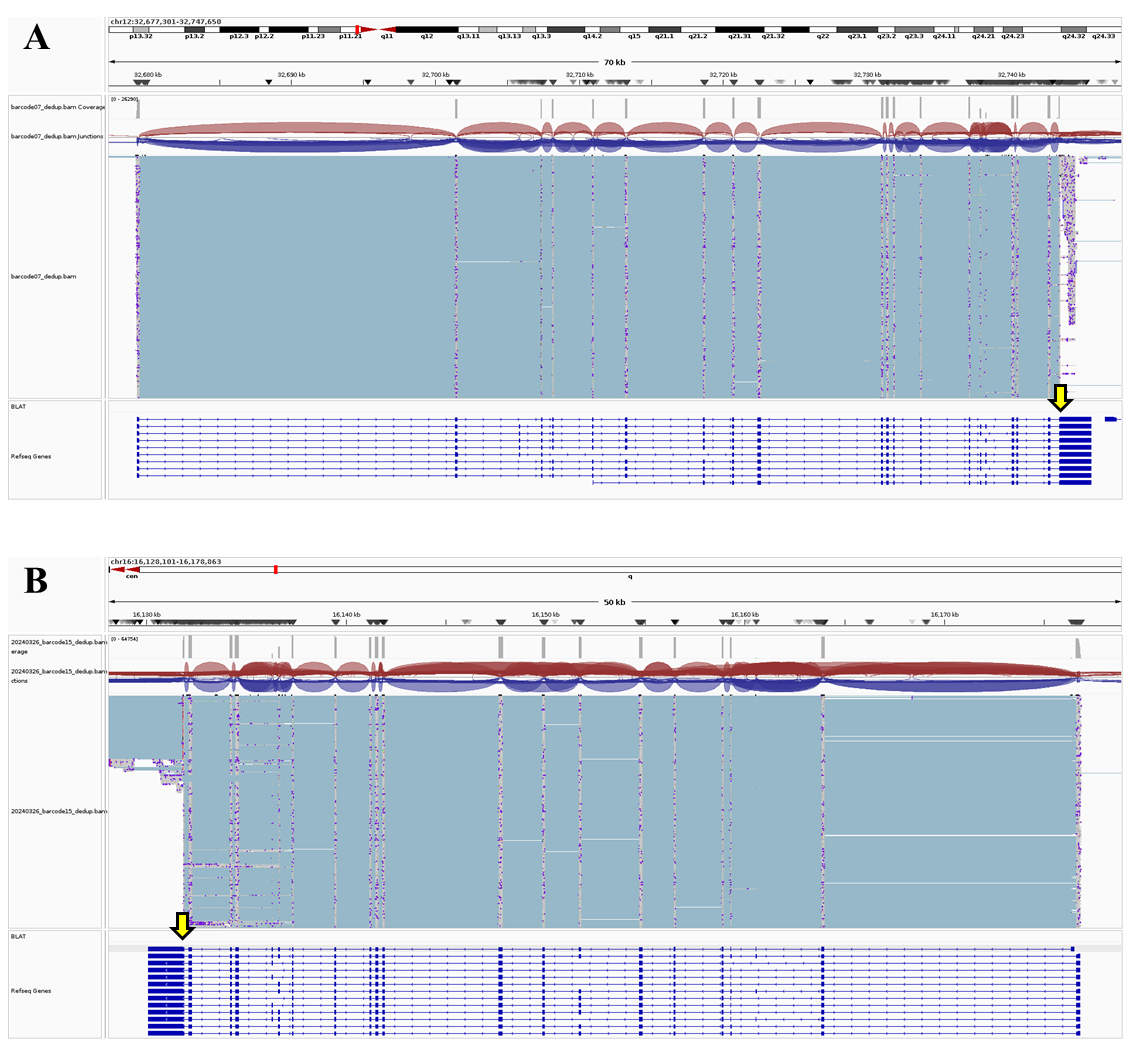

**Supplementary Figure 4:** IGV view of the sequence coverage of the *DNM1L/Dnm1L* in human left ventricle **(A)** and mouse heart **(B)** samples. Yellow arrows indicate the binding position of the primer designed to target exon 21 for amplification.

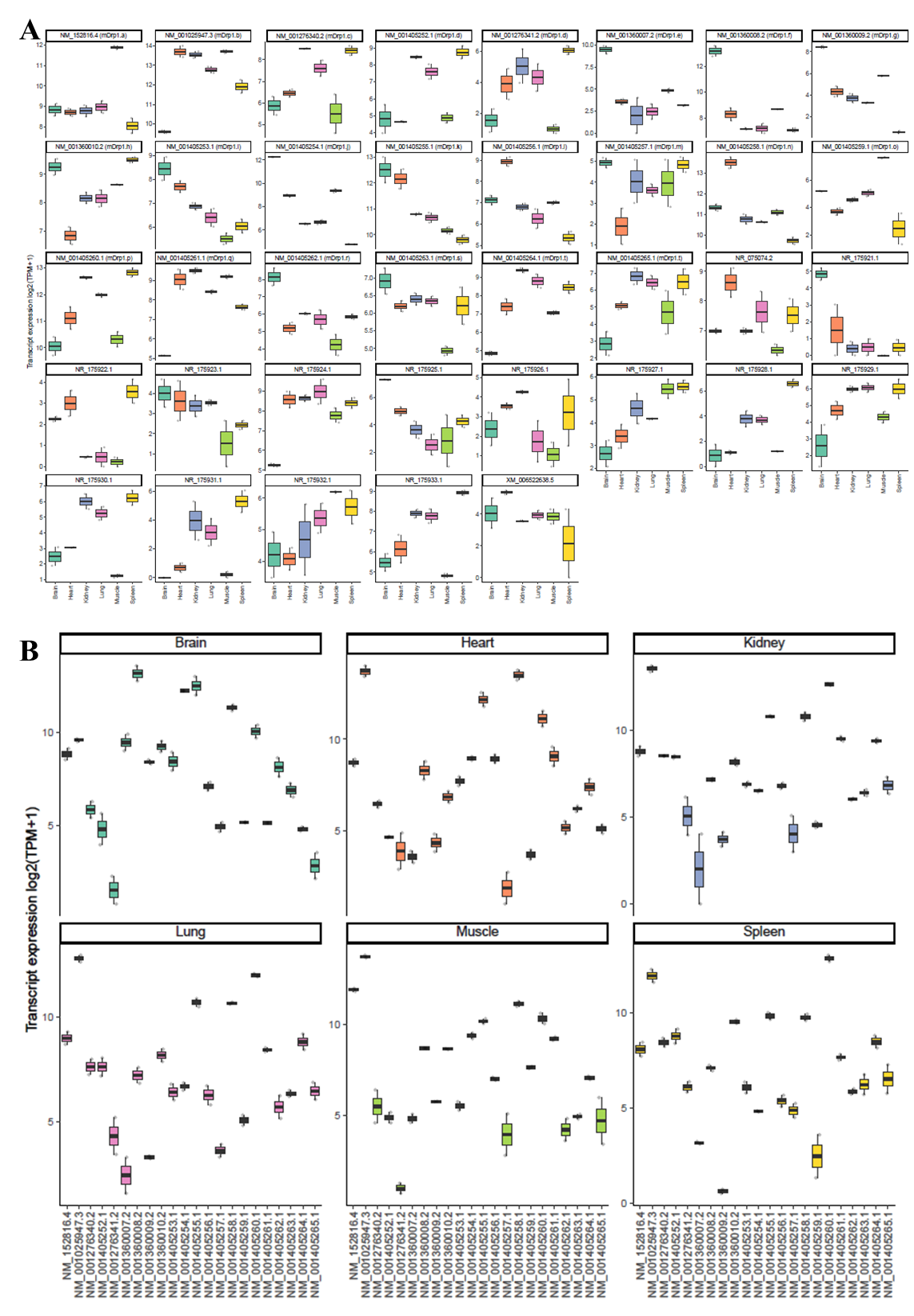

**Supplementary Figure 5: Expression of *Dnm1l* transcript variants in mouse tissues. (A)** Boxplots showing expression levels [log2(TPM+1)] of individual *Dnm1l* transcripts across tissue samples. **(B)** Boxplots showing expression levels [log2(TPM+1)] of individual *Dnm1l* transcripts in different tissue types.
